# Transcriptome profiling of human pluripotent stem cell-derived cerebellar organoids reveals faster commitment under dynamic conditions

**DOI:** 10.1101/2021.01.27.428468

**Authors:** Teresa P. Silva, Rui Sousa-Luís, Tiago G. Fernandes, Evguenia P. Bekman, Carlos A. V. Rodrigues, Sandra H. Vaz, Leonilde M. Moreira, Yas Hashimura, Sunghoon Jung, Brian Lee, Maria Carmo-Fonseca, Joaquim M. S. Cabral

**Affiliations:** iBB – Institute for Bioengineering and Biosciences and Department of Bioengineering, Instituto Superior Técnico, Universidade de Lisboa, Portugal; Instituto de Medicina Molecular João Lobo Antunes, Faculdade de Medicina, Universidade de Lisboa, Portugal; Instituto de Farmacologia e Neurociências, Faculdade de Medicina da Universidade de Lisboa, Portugal; PBS Biotech, CA, USA

**Keywords:** organoids, human pluripotent stem cells, cerebellum, large-scale production, dynamic conditions

## Abstract

Human induced pluripotent stem cells (iPSCs) have great potential for disease modeling. However, generating iPSC-derived models to study brain diseases remains a challenge. In particular, the ability to recapitulate cerebellar development *in vitro* is still limited. We presented a reproducible and scalable production of cerebellar organoids by using the novel Vertical-Wheel single-use bioreactors, in which functional cerebellar neurons were obtained. Here, we evaluate the global gene expression profiles by RNA sequencing (RNA-seq) across cerebellar differentiation, demonstrating a faster cerebellar commitment in this novel dynamic differentiation protocol. Furthermore, transcriptomic profiles suggest a significant enrichment of extracellular matrix (ECM) in dynamic-derived cerebellar organoids, which can better mimic the neural microenvironment and support a consistent neuronal network. Thus, an efficient generation of organoids with cerebellar identity was achieved for the first time in a continuous process using a dynamic system without the need of organoids encapsulation in ECM-based hydrogels, allowing the possibility of large-scale production and application in high-throughput processes. The presence of factors that favors angiogenesis onset was also detected in dynamic condition, which can enhance functional maturation of cerebellar organoids. We anticipate that large-scale production of cerebellar organoids may help developing models for drug screening, toxicological tests and studying pathological pathways involved in cerebellar degeneration.

## Introduction

The human brain represents a complex structure formed by a great diversity of neurons, astrocytes, oligodendrocytes and microglia. Endogenous human brain tissue is not easily available for studying neurodevelopment and neurodegenerative diseases, and it is a subject of ethical concerns. Since the discovery of human pluripotent stem cells (PSCs), including embryonic and induced pluripotent stem cells (ESCs and iPSCs) (Takahashi et al., 2007; Thomson, 1998), distinct approaches have emerged to differentiate them into a variety of glial and neuronal cell types to model human development and neurodegenerative disorders (Ishida et al., 2016; Kim et al., 2019; Liu and Zhang, 2010; Ponroy Bally et al., 2020). However, the reproducible differentiation of a desired neuronal type for disease modeling under defined conditions remains a challenge, aggravated by culture and cell line variability. Engineered 3D cultures resembling complex brain regions, usually called organoids, have been reported (Bagley et al., 2017; Lancaster et al., 2013; Matsumoto et al., 2020; Muguruma et al., 2015; Qian et al., 2018; Xiang et al., 2019). To promote PSC aggregation and generate controlled size and shape PSC aggregates, scaffold-free approaches have been used, including low-cell-adhesion 96-well culture plates (Bagley et al., 2017; Lancaster et al., 2013; Matsumoto et al., 2020; Muguruma et al., 2015; Xiang et al., 2019), conical tubes (Qian et al., 2018) or microwell culture plates (Silva et al., 2020a), which are difficult to adapt for large scale production. Furthermore, it is essential to produce organoids large enough to recapitulate tissue morphogenesis and cellular organization without the limitation of oxygen, nutrients and morphogen diffusion to the cells (McMurtrey, 2016). To help address this critical issue, cerebral organoids were already cultured in dynamic systems. Usually, reported protocols rely on the initial neural commitment of PSCs in static conditions, followed by encapsulation of the organoids in extracellular matrix (ECM)-based hydrogels and their transfer to dynamic culture conditions (Lancaster et al., 2012; Qian et al., 2018). Such approach, however, may limit the potential scale-up of organoid production, which is important for drug screening applications.

We described a new approach for the reproducible and scalable generation of organoids that adopt cerebellar identity and further mature into cerebellar neurons under chemically defined and feeder-free 3D dynamic conditions. Indeed, morphological structures similar to human embryonic cerebellum were firstly generated by Muguruma and colleagues in static conditions (Muguruma et al., 2015). However, these organoids only recapitulate the embryonic structure of cerebellar tissue and the differentiation of functional cerebellar neurons was only achieved in 2D culture after organoid dissociation, using either co-culturing with animal (Muguruma et al., 2015; Tao et al., 2010) and human-derived feeder cells (Wang et al., 2015), or in a co-culture free system (Silva et al., 2020a). By using the novel single-use Vertical-Wheel™ bioreactors (VWBRs, PBS Biotech), we are able to mimic later stages of human cerebellar development *in vitro*, by easily generating high numbers of human iPSC-derived aggregates and efficiently differentiating them into mature cerebellar organoids, which contain diverse types of cerebellar neurons including Purkinje cells and granule cells. VWBRs were already successfully used for human iPSC (Borys et al., 2020; Nogueira et al., 2019; Rodrigues et al., 2018) and mesenchymal stem cell expansion (Sousa et al., 2015; de Sousa Pinto et al., 2019). The VWBRs combine a large vertical impeller and a U-shaped bottom to provide a more homogeneous shear distribution inside the bioreactor, allowing a gentle and uniform mixing and particle suspension with reduced power input and agitation speeds (Croughan et al., 2016). Here, expression profiles from RNA sequencing (RNA-seq) of cerebellar organoids generated under static or dynamic conditions were evaluated, revealing a more efficient cerebellar commitment in the latter protocol. Furthermore, RNA-seq data analysis suggests a significant enrichment of ECM in dynamic conditions, avoiding the encapsulation of organoids in ECM-based hydrogels and thus facilitating the large-scale production and their applicability in high-throughput processes. The presence of a microenvironment that favors angiogenesis onset was also observed in dynamic conditions, which can support a more complex environment sustaining a functional maturation of cerebellar organoids.

## Materials and Methods

### Maintenance of human iPSCs

In this study, three distinct human iPSC lines, F002.1A.13 (Silva et al., 2020a), Gibco Human Episomal iPSC line (iPSC6.2, Thermo Fisher Scientific) (Burridge et al., 2011) and iPS-DF6-9-9T.B (WiCell Bank)(Junying et al., 2009) were used. All human iPSCs were cultured on Matrigel (Corning)-coated 6-well plates with mTeSR™1 medium (StemCell Technologies). Full-volume medium replacement with mTESR1 was performed daily. Cells were passaged when the colonies covered approximately 85% of the surface area of the culture dish at a split ration of 1:3, using 0.5mM EDTA dissociation buffer (Thermo Fisher Scientific)(Beers et al., 2012). Two to three passages were performed before starting the differentiation protocol.

### Generation, differentiation and maturation of human iPSC-derived aggregates using Vertical-Wheel Bioreactors

In this work, PBS MINI 0.1 MAG VWBRs (PBS Biotech, USA) were used, which hold a maximum volume of 100mL. The working volume selected was 60mL, which allows a complete covering of the impeller wheel with culture medium. The protocols for seeding, operation of the vessel, and cerebellar differentiation of human iPSCs are described in detail elsewhere (Silva et al., 2020a; Silva et al., 2020b). Briefly, for single-cell seeding, cells grown in a culture dish were incubated with ROCK inhibitor (Y-27632, 10μM, StemCell Technologies) for 1 h at 37°C prior to harvesting with Accutase. Then, cells were treated with accutase (Sigma) for 7 min at 37°C. After dissociation, single-cells were seeded in the bioreactor at a density of 250 000 cells/mL in 60mL of mTeSR™1 supplemented with 10μM Y-27632. To promote cell aggregation an agitation speed of 27 rpm was used. After 24 hours, 80% of the medium was replaced and aggregates were maintained in mTeSR™1 without Y-27632 for another 24 hours at an agitation speed of 25 rpm. From day 2 to day 21 after seeding, gfCDM was used as basal medium for the differentiation (Muguruma et al., 2015; Silva et al., 2020a). Recombinant human basic FGF (FGF2, 50ng/ml, PeproTech) and SB431542 (10μM, Sigma) were added to culture on day 2. Full-volume medium replacement with gfCDM (supplemented with insulin, FGF2 and SB431542) was performed on day 5, letting the organoids settle at the bottom of the bioreactor. On day 7, the agitation speed was changed to 30 rpm, medium was fully replaced and two-thirds of initial amounts of FGF2 and SB were added. Recombinant human FGF19 (100ng/ml, PeproTech) was added to culture on day 14 post-seeding, and full-volume replacement was performed on day 18. From day 21, the aggregates were cultured in Neurobasal medium (Thermo Fisher Scientific) supplemented with GlutaMax I (Thermo Fisher Scientific), N2 supplement (Thermo Fisher Scientific), and 50U/ml penicillin/50μg/ml streptomycin (PS, Thermo Fisher Scientific). Full-volume replacement was performed every 7 days. Recombinant human SDF1 (300ng/ml, PeproTech) was added to culture on day 28. After 35 days of differentiation, neuronal maturation was promoted by using BrainPhys Neuronal Medium (StemCell Technologies), supplemented with NeuroCult SM1 Neuronal Supplement (StemCell Technologies), N2 Supplement-A (StemCell Technologies), Recombinant Human Brain Derived Neurotrophic Factor (BDNF, PeproTech, 20ng/mL), Recombinant Human Glial-Derived Neurotrophic Factor (GDNF, PeproTech, 20ng/mL), dibutyryl cAMP (1mM, Sigma), and ascorbic acid (200nM, Sigma). One-third of total volume was replaced every 3 days.

### Aggregate size and biomass analysis

To monitor aggregate sizes throughout time in culture, several images were acquired at different time points using a Leica DMI 3000B microscope with a Nikon DXM 1200F digital camera. The aggregate area was measured using ImageJ Software. Considering the aggregates as spheroids, diameters were calculated based on determined area according to the equation: 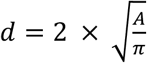 in which *d* represents the diameter and *A* represents the area. To analyze the volume of biomass, the volume was calculated based on the determined average diameter according to the equation: 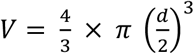. The total increase of biomass was calculated multiplying the volume of an aggregate in average by the total number of organoids and normalized to the biomass measured on day 1. The number of aggregates were manually quantified using 1mL-sample collected from the bioreactor.

### Expression profiling with RNA sequencing. 1) Sample collection and RNA extraction

F002.1A.13 iPSC line-derived aggregates were collected at different time-points of cerebellar differentiation from VWBRs and static conditions. For RNA extraction, aggregates were dissociated with accutase at 37°C for 7 min. After enzymatic neutralization, the cell pellet was washed with phosphate buffered saline (PBS, 0.1M) and then stored at −80°C. Total RNA was extracted from samples using High Pure RNA Isolation Kit (Roche, Cat. 11828665001), according to the manufacturer instructions. **2) RNA-seq sample preparation and sequencing.** RNA libraries were prepared for sequencing using Lexogen QuantSeq 3’mRNA-Seq Library Prep Kit FWD for Illumina using standard protocols. Briefly, 500ng of total RNA were primed with the oligo dT primer containing Illumina-compatible linker sequences. After first strand synthesis, the RNA was removed, and second strand synthesized with Illumina-compatible random primers. After magnetic bead-based purification, the libraries were PCR amplified introducing the sequences required for cluster generation. Sequencing was performed using NextSeq (75 cycles protocol) platforms. Base calling of samples processed in NextSeq Sequencer was performed with the Real-Time Analysis (RTA) v2. **3) Transcriptome analyses.** Quality control of raw Illumina reads was performed using FastQC v0.11.5 tool. TrimGalore v0.4.4 was employed to trim read adaptors in paired-end mode, removing reads with less than 10 bases and/or low-quality ends (20 Phred score cut-off). The resultant reads were aligned against the reference human genome (GRCh38) using STAR v2.7.0 software, requiring uniquely mapped reads (--outFilterMultimapNmax 1) and minimum alignment score (--outFilterScoreMin) of 10. BAM files with aligned reads were run through featureCounts v2.2.6 (strandSpecific = 1) to produce estimated gene expression values, which were then gathered in a non-normalized count matrix. Normalization of gene’s read counts comparison between static and dynamic protocols as well as different timepoints were done using DESeq2 v1.28.1 rlog function. PCA plot was performed using plotPCA function from the same package. Heatmaps were built using pheatmap package. Significant differentially expressed genes were detected with DESeq2 package and R v4.0.2. A double cut-off of 0.05 for adjusted p-value and 2 for | log2(FoldChange) | was applied over DESeq2’s own two-sided statistical test results. Gene set enrichment analyses was performed with topGO v2.40.0, which allowed to revel the overrepresented GO terms between conditions. **4) Accession numbers.** RNA-seq data for this study are available through Gene Expression Omnibus (GEO) Accession Number GSE161549

### Quantitative Real-time PCR (qRT-PCR)

Total RNA was extracted at different time-points of cerebellar differentiation using High Pure RNA Isolation Kit (Roche) and converted into complementary cDNA with Transcriptor High Fidelity cDNA Synthesis Kit (Roche). Gene expression was analyzed using SYBR^®^ green chemistry (**Table 1**). All PCR reactions were run in triplicate, using the ViiA™7 RT-PCR System (Applied BioSystems). Quantification was performed by calculating the ΔCt value using GAPDH as a reference and results are shown as mRNA expression levels (2-ΔCt) relative to GAPDH.

**Table 1.**
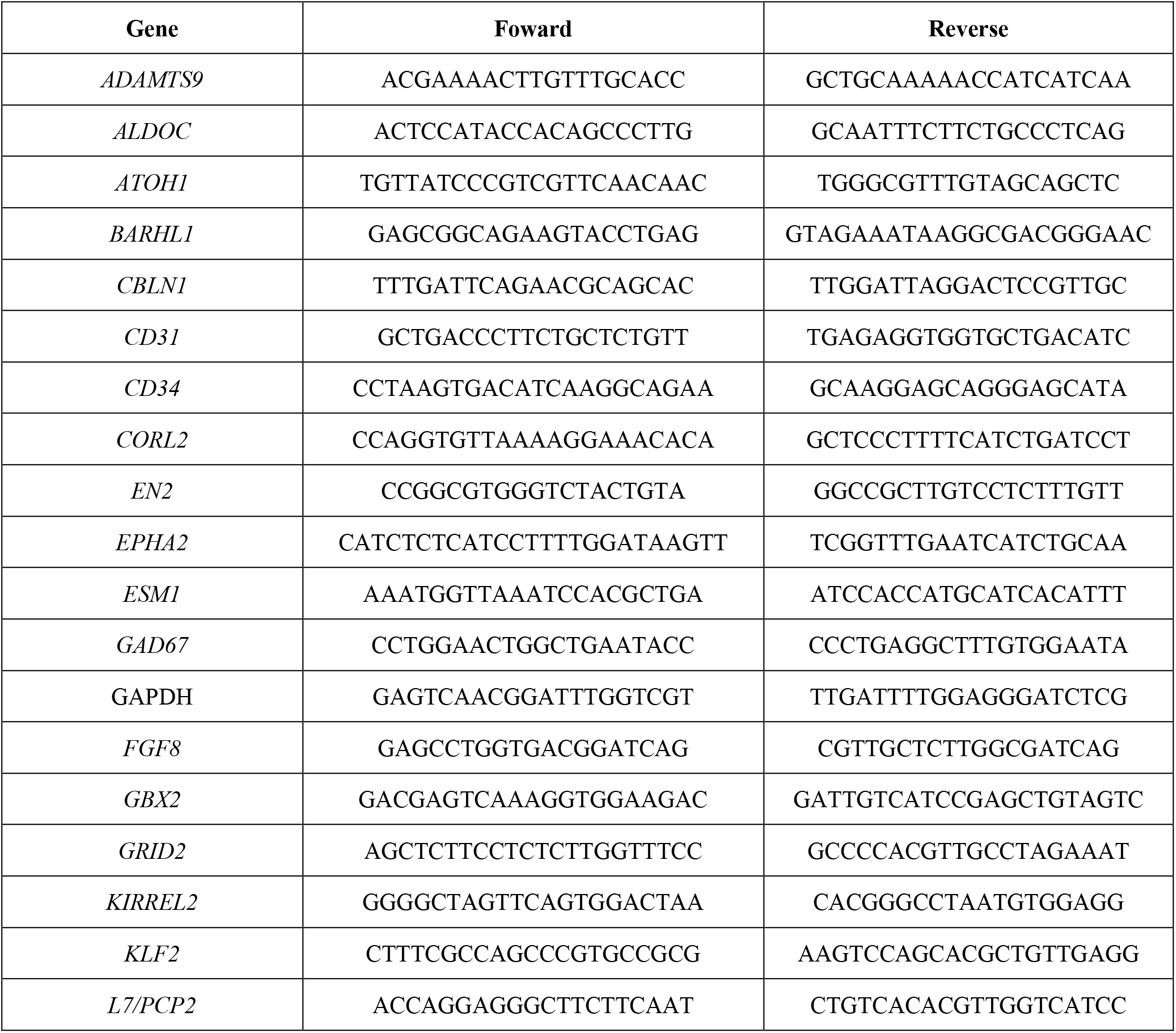
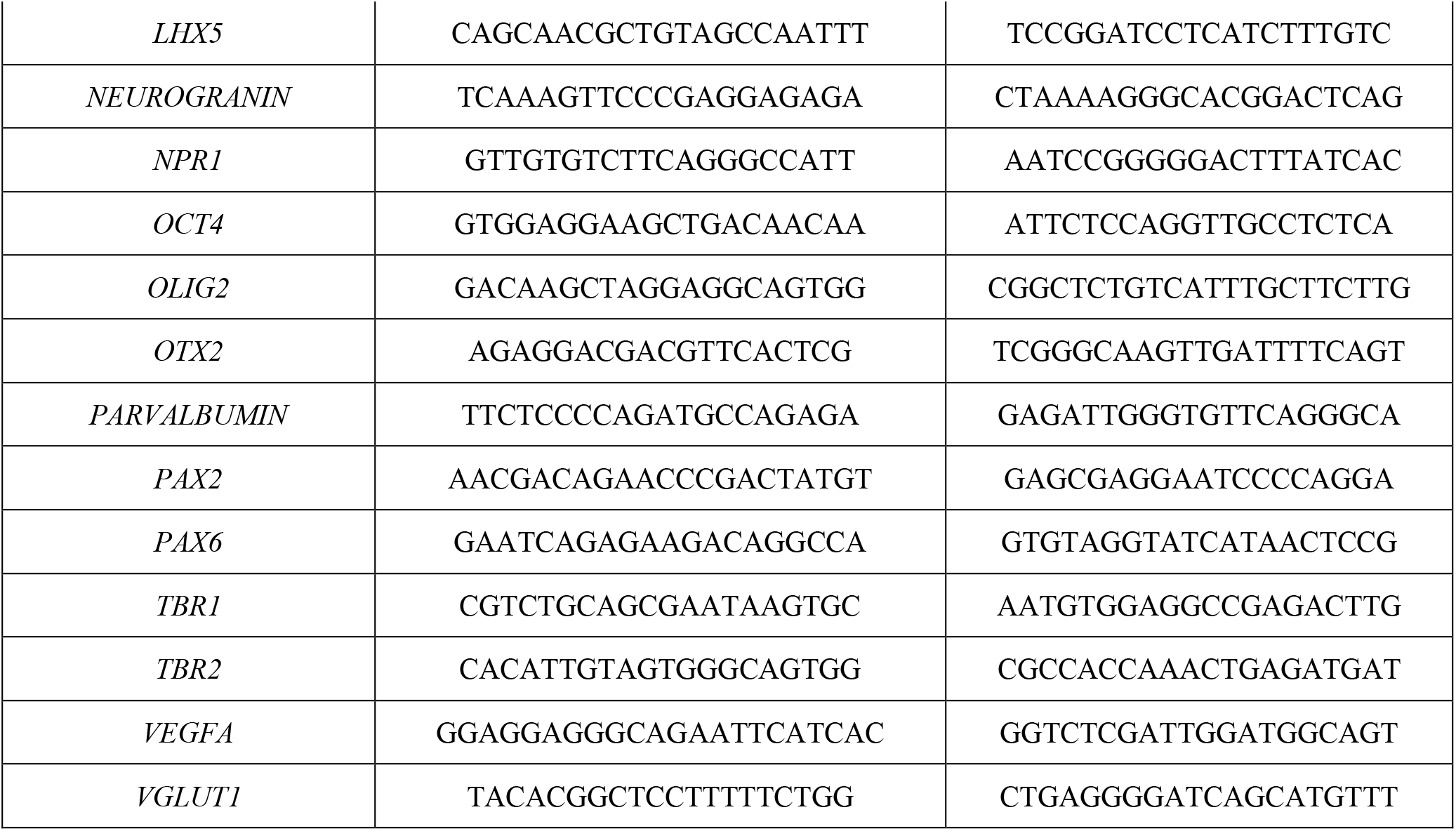
Primers used for qRT-PCR

### Tissue preparation and Immunohistochemistry

Aggregates were fixed in 4% paraformaldehyde (PFA, Sigma) for 45 min at 4°C followed by washing in PBS and overnight incubation in 15% (v/v) sucrose at 4°C. Aggregates were embedded in 7.5%/15% (v/v) gelatin/sucrose and frozen in isopenthane at −80°C. Twelve-μm sections were cut on a cryostat-microtome (Leica CM3050S, Leica Microsystems), collected on Superfrost™ Microscope Slides (Thermo Scientific) and stored at −20°C. For immunostaining, sections were de-gelatinized for 45 min in PBS at 37°C, incubated in 0.1 M Glycine (Millipore) for 10 min at room temperature (RT), permeabilized with 0.1% (v/v) Triton X-100 (Sigma) for 10 min at RT and blocked with 10% (v/v) fetal bovine serum (FBS, Gibco) in TBST (20 mM Tris-HCl pH 8.0, 150 mM NaCl, 0.05 % v/v Tween-20, Sigma) for 30 min at RT. Sections were then incubated overnight at 4°C with the primary antibodies diluted in blocking solution (**Table 2**). Secondary antibodies were added to sections for 30 min (goat anti-mouse or goat anti-rabbit IgG, Alexa Fluor^®^–488 or –546, 1:400 v/v dilution, Molecular Probes) at RT and nuclear counterstaining was performed using 4’,6-diamidino-2-phenylindole (DAPI, 1.5 μg/mL; Sigma). After brief drying, sections were mounted in Mowiol (Sigma). Fluorescence images were acquired with Zeiss LSM 710 Confocal Laser Point-Scanning Microscopes.

**Table 2.**
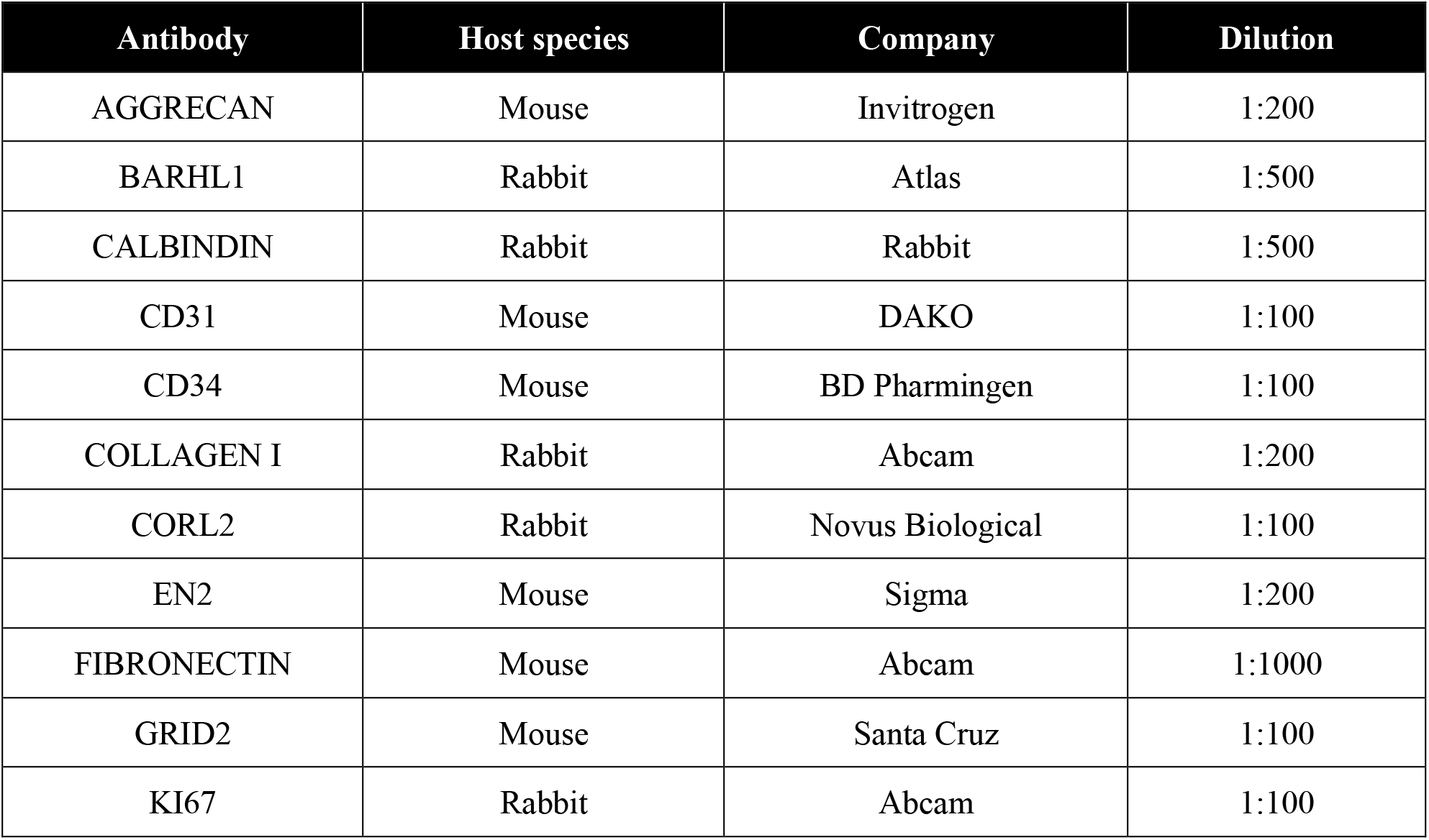
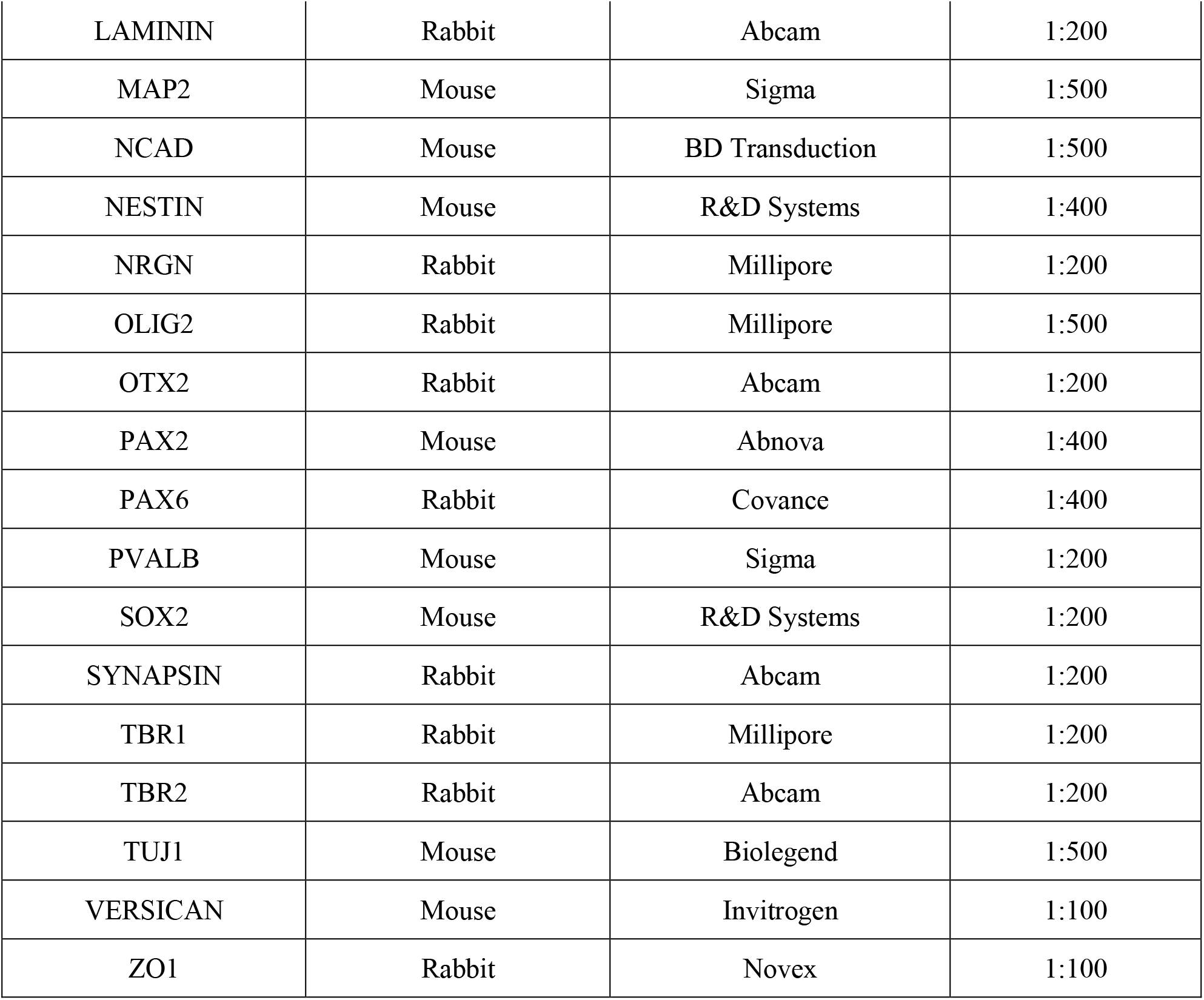
List of primary antibodies and dilutions used for immunostaining

### Single cell calcium imaging

Functional maturation was evaluated by single cell calcium imaging (SCCI) to analyze the intracellular variations of Ca^2+^ following stimulation with 50mM KCl and 100μM histamine (Sigma). 2-5 days before evaluation, aggregates were re-plated on Glass Bottom Cell Culture Dish (Nest) previously coated with poly-L-ornithine (15μg/mL, Sigma) and laminin (20μg/mL, Sigma). At different time points of differentiation, neurons were loaded with Fura-2 AM (5 μM, in normal Krebs solution with the following composition: NaCl (132mM), KCl (4mM), MgCl_2_ (1.4mM), CaCl_2_ (2.5mM), D-(+)-glucose (6mM) and HEPES (10mM) - pH 7.4 adjusted with NaOH - and incubated at 37°C for 45 min. Fura-2 AM loaded cells were sequentially excited both at 340nm and 380nm, for 250ms at each wavelength, using an inverted microscope with epifluorescent optics and equipped with a high speed multiple excitation fluorimetric system (Lambda DG4, with a 175W Xenon arc lamp). The emission fluorescence was recorded at 510nm by a CDD camera. Cells were stimulated using 100μM histamine or high potassium Krebs solution (containing 50mM KCl, isosmotic substitution with NaCl), as reported elsewhere (Rodrigues et al., 2017; Xapelli et al., 2014).

### Patch Electrophysiology

For electrophysiological evaluation, 2-5 days before the analysis aggregates were gently dissociated using accutase (Sigma) and re-plated on coverslips coated with poly-L-ornithine (15μg/mL, Sigma) and Laminin (20μg/mL, Sigma). Whole cell patch-clamp recordings were obtained from generated neurons using an upright microscope (Zeiss Axioskop 2FS) equipped with differential interference contrast optics using a Zeiss AxioCam MRm camera and a 40x IR-Achroplan objective. During recordings, cells were continuously perfused with artificial cerebrospinal fluid containing: 124mM NaCl, 3mM KCl, 1.2mM NaH_2_PO_4_, 25mM NaHCO_3_, 2mM CaCl_2_, 1mM MgSO_4_ and 10mM glucose, which was continuously gassed with 95%O_2_/5% CO_2_. Recordings were performed at room temperature in current-clamp [holding potential (Vh) = −70 mV] with an Axopatch 200B (Axon Instruments) amplifier, as performed in (Felix-Oliveira et al., 2014). Briefly, patch pipettes with 4 to 7 MΩ resistance when filled with an internal solution containing: 125mM K-gluconate, 11mM KCl, 0.1mM CaCl_2_, 2mM MgCl_2_, 1mM EGTA, 10mM HEPES, 2mM MgATP, 0.3mM NaGTP, and 10mM phosphocreatine, pH 7.3, adjusted with NaOH. 280-290 mOsm were used to record action potential activity. Acquired signals were filtered using an in-built, 2-kHz, three-pole Bessel filter, and data were digitized at 5 kHz under control of the pCLAMP 10 software program. The junction potential was not compensated for, and offset potentials were nulled before gigaseal formation. The resting membrane potential was measured immediately upon establishing whole-cell configuration. Firing patterns of cerebellar neurons were determined in current-clamp mode immediately after achieving whole-cell configuration by a series of hyperpolarizing and depolarizing steps of current injection (500ms). Firing potential were also determined through the application of two depolarizing steps of current injection of 10ms, separated by 80ms.

## Results

### Production of size-controlled human iPSC-derived aggregates leads to an efficient neural commitment

The generation of human iPSC-derived organoids starts by cell aggregation to mimic the 3D structure and recapitulate both organization and functionality of human organs. The aggregation process is a critical step to obtain a homogeneous outcome in the efficiency of differentiation with high yield of viable organoids and increased reproducibility of the protocol (Xie et al., 2017). We initiated the protocol by promoting cell aggregation using VWBRs (**Supplementary Fig. 1a**), in which single-cell seeding was performed at 250 000 cells/mL in 60 mL of medium and with an agitation speed of 27 rpm (**Fig. 1a**). After 24 hours, cells were able to efficiently aggregate (**Supplementary Fig. 1b**) and homogeneously shaped aggregates were obtained (day 1, **Fig. 1b**). Distribution of diameters showed that aggregate size continues to increase from day 1 to 5, achieving similar size between different iPSC lines. Furthermore, homogeneous size and shape aggregates were observed when differentiation was initiated at day 2 and their spheroid-like structure was maintained during this initial neural commitment and differentiation (**Fig. 1b**). Further analysis of aggregate diameters revealed that organoids were able to grow in size until day 35 (**Fig. 1c**), with exception of iPSC6.2 cell line that presented no differences between day 21 and 35. A large number of iPSC-derived organoids can be generated with the bioreactor using a straightforward methodology, achieving about 350 ± 52 (mean ± SEM) aggregates/mL 24 hours after seeding. This number decreased at day 2 but remained constant until the end of the cerebellar commitment, achieving 3201 ± 123 (mean ± SEM) organoids in a VWBR with a working volume of 60 mL (**Fig. 1d** and **Supplementary Fig. 1c**). Probably, the merging of individual aggregates was promoted by the decrease in the agitation speed from 27 to 25 rpm, since biomass analysis demonstrated that the total volume of biomass increased up to ~4-fold until day 7, achieving a ~6-fold increase on day 35 of differentiation (**Fig. 1e** and **Supplementary Fig. 1d**).

**Figure 1.**
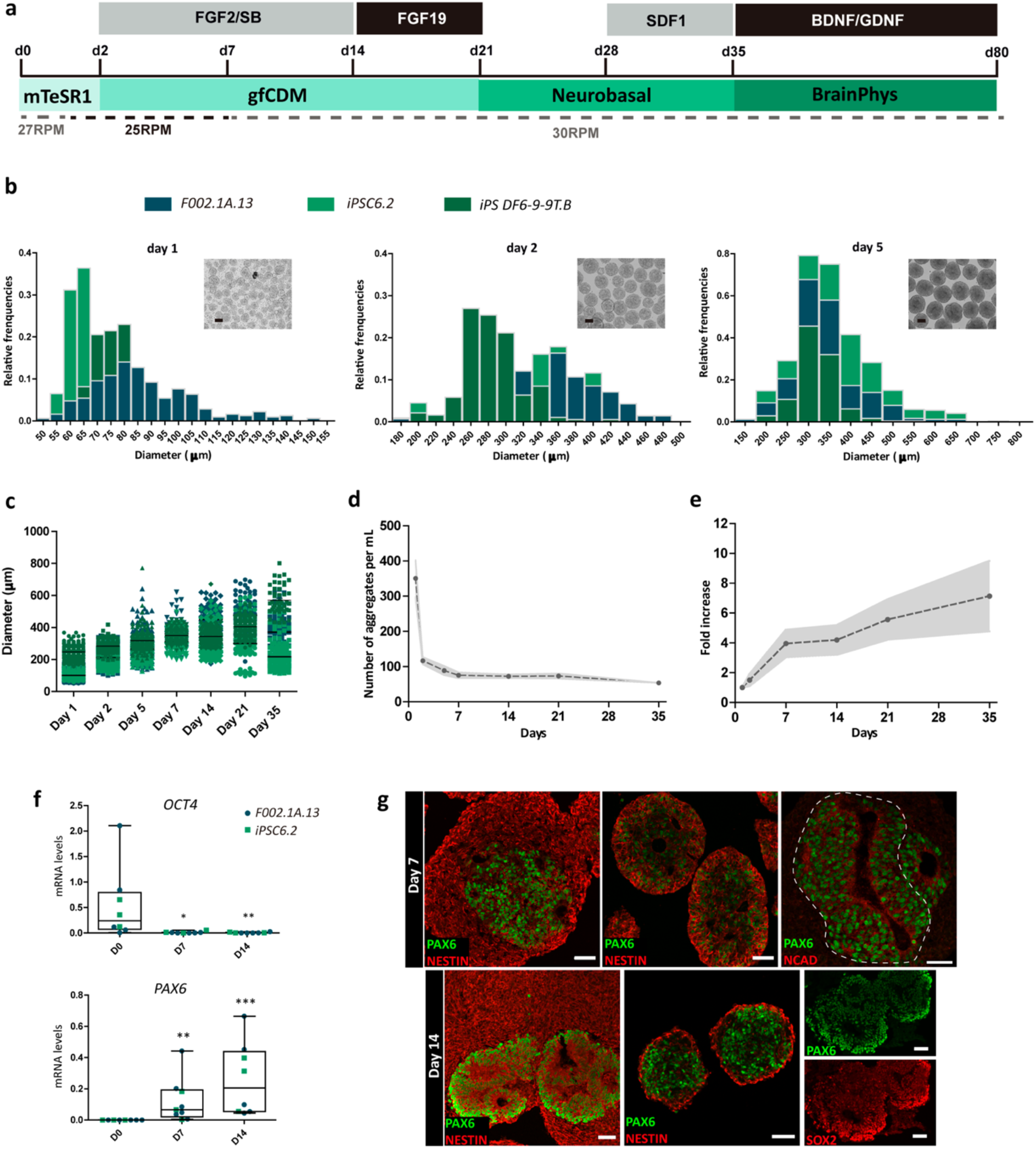
Neural commitment in human iPSC-derived aggregates using VWBRs. **a,** Schematic for cerebellar differentiation of human iPSC-derived aggregates using VWBR (see also Methods for details). **b,** Distribution of floating aggregates diameters and representative bright field photomicrographs at days 1, 2 and 5 of differentiation, demonstrating that homogeneous size and shape aggregates were obtained from different human iPSC lines using VWBRs. **c,** Diameter of iPSC-derived aggregates along the cerebellar differentiation using different human iPSC lines. **d,** Number of iPSC-derived aggregates from day 1 to day 35 of cerebellar differentiation. Points show the mean of 3 independent differentiation experiments using 3 different iPSC lines and the area fill represents SEM (see also Supplementary Fig. 1 for more details). **e,** Total volume of biomass relative to day 1. Points show the mean of 3 independent differentiation experiments using 3 different iPSC lines and the area fill represents SEM (see also Supplementary Fig. 1 for more details). **f,** qRT-PCR analyses of aggregates derived from different iPSC lines at the indicated time-points. Box-plots diagrams depict mRNA expression levels (2^-ΔCT^) relative to *GAPDH*. Each color and shape-coded dot represents data from one experiment. One-way ANOVA (Kruskal-Wallis test), *p<0.05, **p<0.01, ***p<0.001; error bars represent SEM. **g,** Immunofluorescence for NESTIN, PAX6, NCAD and SOX2 on days 7 and 14 of differentiation. Scale bars, 50μm.

To induce neural commitment, aggregates were cultured in neural medium supplemented with SB431542, fibroblast growth factor 2 (FGF2) and insulin, which are sufficient to promote neural differentiation with moderate caudalization of the neuroepithelium that is essential to mid-hindbrain patterning. Subsequently, fibroblast growth factor 19 (FGF19) and stromal cell-derived factor 1 (SDF1) were sequentially introduced in the culture to promote tissue polarity and generate different cerebellar progenitors. The pluripotency and self-renewal transcript *OCT4* was detected in human iPSCs on day 0 and upon neural induction its mRNA levels were down-regulated to almost undetectable levels by day 7 (**Fig. 1f**). Consistent with this significant downregulation of the pluripotency gene, the mRNA levels of *PAX6*, which is a transcription factor driving neurogenesis and important for neural stem cell proliferation (Sansom et al., 2009; Thakurela et al., 2016), significantly increased, showing that an efficient neural commitment was already reached by day 7 (**Fig. 1f**). Immunofluorescence analysis further supports that an efficient neural commitment of the iPSC-derived aggregates is achieved by day 7 of differentiation, with most of cells within the aggregates expressing the neural progenitor marker NESTIN (**Fig. 1g**). The cryosections of organoids also revealed many structures reminiscent of the neural tube, expressing the neural markers PAX6 and SOX2 (**Fig. 1g**). Furthermore, these neural tube-like structures showed apico-basal polarity with PAX6 and SOX2 co-stained progenitors found at the luminal (apical) side marked by strong expression of apical marker N-cadherin (NCAD, **Fig. 1g**).

### Induction of cerebellar identity in 3D dynamic culture conditions

During human development, the territory that gives rise to the cerebellum is located in one of the hindbrain segments, which comprises the most anterior zone of the hindbrain caudally to the mid-hindbrain boundary (MHB), the isthmic organizer (IsO)(Zervas et al., 2004), and is established by differential expression of several transcription factors (Watson et al., 2015). The expression levels of *OTX2* and *GBX2* transcripts (**Fig. 2a**), that define the molecular limits of MHB, were significantly upregulated by day 7, together with those of *FGF8, EN2* and *PAX2*, which are crucial transcription factors involved in IsO specification (Chi et al., 2003; Joyner, 1996; Martinez et al., 1999) (**Fig. 2a**). Immunostaining analysis of organoids supports the efficient self-formation of IsO tissue, staining for OTX2 and EN2 by day 14 (**Fig. 2b, *i* and *ii***). Therefore, signals from this organizer center, including FGF8 (**Fig. 2a**), were able to stimulate the generation of cerebellar territory (Sasai, 2013), supported by expression of PAX2 (**Fig. 2b, *iii***), a marker for GABAergic interneuron precursors in the developing cerebellum (Maricich and Herrup, 1999), and BARHL1 (**Fig. 2b, *iv***), a marker for glutamatergic progenitors during cerebellar development (Li, 2004). The validation of efficient neuronal differentiation was performed by transcriptomic analysis and supported by gene ontology (GO) analysis for upregulated genes on day 14 compared with day 0 (**Supplementary Fig. 1e-f**). The differential expression analysis between days 14 and 0 demonstrated an enrichment of genes on day 14 that are crucial for metencephalon development, the tissue that differentiates into pons and cerebellum, and further cerebellar differentiation (**Fig. 2c**), including *EN2, PAX2* and *BARLH1*, supporting an effective cerebellar commitment in human iPSC-derived organoids already by day 14. By day 21, human iPSC-derived organoids displayed continuous neuroepithelial layer with PAX6^+^ neural progenitors on the surface with few TUJ1^+^ newborn neurons within the organoid (**Fig. 2d, *i***). Furthermore, consistent with cerebellar identity, BARHL1^+^ progenitors self-organized into continuous layers (**Fig. 2d, *ii***) basally located to proliferating SOX2^+^ progenitors (**Fig. 2d, *iii***). Other types of cerebellar progenitors were found by day 21. These cells were positive for OLIG2, a marker of neurogenic progenitors of Purkinje cells (Seto et al., 2014), and were dispersed within the organoids (**Fig. 2d, *iv***). On day 35, after treatment with chemokine that regulates cerebellar migration (Bagri et al., 2002), SDF1, GO analysis of significantly upregulated genes (compared to day 0) revealed enrichment of neurological processes, including neuron migration and synaptic processes (**Supplementary Fig. 1g-h**), supporting the reorganization of the neuroepithelium and initiation of neuronal maturation. Human-iPSC derived organoids were polarized by day 35, with staining for ZO1 (**Fig. 2e, *i***), an apical marker, on the presumably apical side of the tissue. Two different structures can be observed, neural tube-like structures enclosed within the organoid with apical side towards the lumen (**Fig. 2e, *i*,** asterisk), and a continuous neuroepithelium that contains the apical side on the outer surface of the organoid (**Fig. 2e, *i*,** arrowheads), probably resulting from merging of neural tube-like structures promoted by SDF1 addition (Muguruma et al., 2015). In both cases the internal layer organization maintained the same pattern, with a layer of proliferating progenitors SOX2^+^ (**Fig. 2e, *ii***) and PAX6^+^ always found on the apical side (**Fig. 2e, *iii***), and TUJ1^+^ neuronal cells disposed basally to PAX6^+^ neural progenitors within the organoids (**Fig. 2e, *iii***). Consistent with this organization, post-mitotic BARHL1^+^ glutamatergic cerebellar progenitors were located basally within the organoid (**Fig. 2e, *iv***). Other cerebellar progenitor populations were detected by day 35, including cells expressing PAX2 (**Fig. 2e, *v***) and few dispersed cells expressing CORL2 (**Fig. 2e, *vi***), a marker for precursors of Purkinje cells (Nakatani et al., 2014). The differential expression analysis between days 35 and 0 supported an efficient cerebellar commitment and differentiation (**Fig. 2f**), which was validated by qRT-PCR, with significant expression of transcripts encoding specific markers for different types of cerebellar progenitors (**Fig. 2g**). These markers included GABAergic cerebellar progenitors: *KIRREL2* (essential regulator of GABAergic neuron development(Mizuhara et al., 2010)), *PAX2, OLIG2* and *CORL2*, and glutamatergic precursors: *PAX6, ATOH*1 (essential for genesis of granule cells (Ben-Arie et al., 1997) and *BARLH1*.

**Figure 2.**
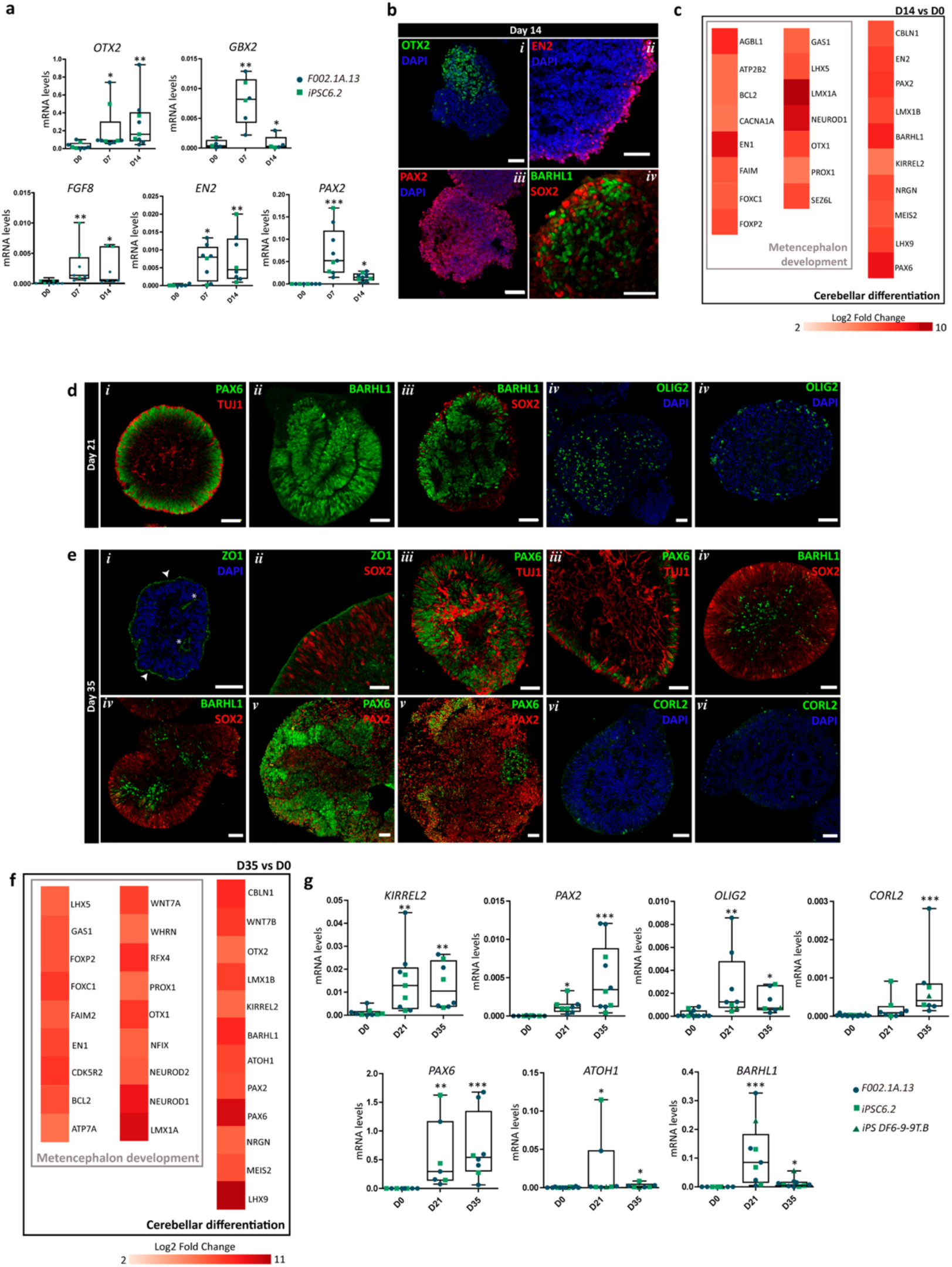
Efficient cerebellar commitment and differentiation in human iPSC-derived aggregates using VWBRs. **a,** qRT-PCR analysis of aggregates derived from different human iPSC lines at the indicated time-points. Box-plots diagrams depict mRNA expression levels (2^-ΔCT^) relative to *GAPDH*. Each color and shape-coded dot represents data from one differentiation experiment. One-way ANOVA (Kruskal-Wallis test), *p<0.05, **p<0.01, ***p<0.001; error bars represent SEM. **b,** Immunofluorescence analysis on day 14 of differentiation for OTX2, EN2, PAX2, BARHL1 and SOX2. Scale bars, 50μm. **c,** Heatmap highlighting genes differentially expressed (Log2 FC > 2 and adjusted p-value < 0.05) related with metencephalon development and cerebellar differentiation for day 14 vs day 0 of differentiation. **d and e,** Immunostaining analysis on days 21 and 35 of differentiation for the indicated markers. Scale bars, 50μm. **f,** Heatmap highlighting genes differentially expressed (Log2 FC > 2 and adjusted p-value < 0.05) related with metencephalon development and cerebellar differentiation for day 35 vs day 0 of differentiation. **g,** qRT-PCR analysis of 3D cultures derived from different human iPSC lines at the indicated time-points. Box-plots diagrams depict mRNA expression levels (2^-ΔCT^) relative to *GAPDH*. Each color and shape-coded dot represents data from one differentiation experiment. One-way ANOVA (Kruskal-Wallis test), *p<0.05, **p<0.01, ***p<0.001; error bars represent SEM.

### Identification of different functional cerebellar neurons in dynamic culture of cerebellar organoids

Further maturation of cerebellar organoids was promoted by culturing them in BrainPhys™ medium supplemented with BDNF and GDNF. Gene expression analysis showed, by day 56, the expression of *GAD67* and *VGLUT1* transcripts (**Supplementary Fig. 2a**), which are present in GABAergic and glutamatergic neurons, respectively, and this was maintained until day 80, suggesting the presence of these two major neuronal subtypes in the organoids. Immunostaining analysis further revealed the presence of specific markers for distinct types of cerebellar neurons. The major glutamatergic cerebellar neurons, the granule cells, were detected by co-staining for PAX6 and MAP2 (**Fig. 3a**). Interestingly, immunofluorescence using antibodies to PAX6, expressed early in neural progenitors and later in cerebellar granule cells (Swanson et al., 2005), and MAP2, a neuron-specific microtubule associated protein, revealed a robust organization within organoids. A dense layer of PAX6^+^/MAP2^−^ precursors was detected, whereas MAP2^+^ fibers were distinguished basally to the progenitor’s layer by day 56 of differentiation (**Fig. 3a, *i***), and the co-localization of PAX6^+^ cells with MAP2^+^ neuronal network (**Fig. 3a, *ii***), indicated the presence of mature granule cells. Along the maturation, the initially large PAX6^+^ neuroepithelium became smaller while, simultaneously, the MAP2^+^ region was extended (**Fig. 3a**). In some organoids a mature neuronal network was formed, without the presence of neural progenitors (**Fig. 3a, *iv***), whereas in other cases a niche of PAX6^+^ progenitors remained until day 80 (**Fig. 3a, *v***). Also, other types of glutamatergic cerebellar cells seem to be found, including unipolar brush cells staining for TBR2 and MAP2 (**Fig. 3b**) and deep cerebellar projection neurons expressing TBR1 and MAP2 (**Fig. 3c**). Purkinje cells, the class of GABAergic neurons within the cerebellum with a most elaborated dendritic arbor, expressing the calcium-binding protein calbindin (CALB), were detected within a compact meshwork along the surface of organoids (**Fig. 3d**). Some of the CALB^+^ cells were also positive for Purkinje cell-specific glutamate receptor GRID2 (**Fig. 3e**), indicating the presence of mature Purkinje cells. Within the neuronal network, cells expressing parvalbumin (PVALB) were also found, and their non-overlapping expression with CALB identifies them as GABAergic interneurons (**Fig. 3f**). Golgi cells, another GABAergic cerebellar cell type, were also detected by staining for neurogranin (NRGN) and PAX2 (**Fig. 3g**). In agreement with immunofluorescence analysis, the quantification of transcripts encoding for markers of distinct cerebellar neurons also demonstrated their robust expression during the maturation protocol (**Fig. 3h**). Significant levels of mRNA encoding for specific markers of Purkinje cells were detected by qRT-PCR analysis (**Fig. 3i**), including *L7/PCP2* (Purkinje cell protein 2), *GRID2, CBLN1* (Cerebellin 1 Precursor), *LHX5* (LIM-homeodomain transcription factor) and *ALDOC* (aldolase C, a brain type isozyme of a glycolysis enzyme).

**Figure 3.**
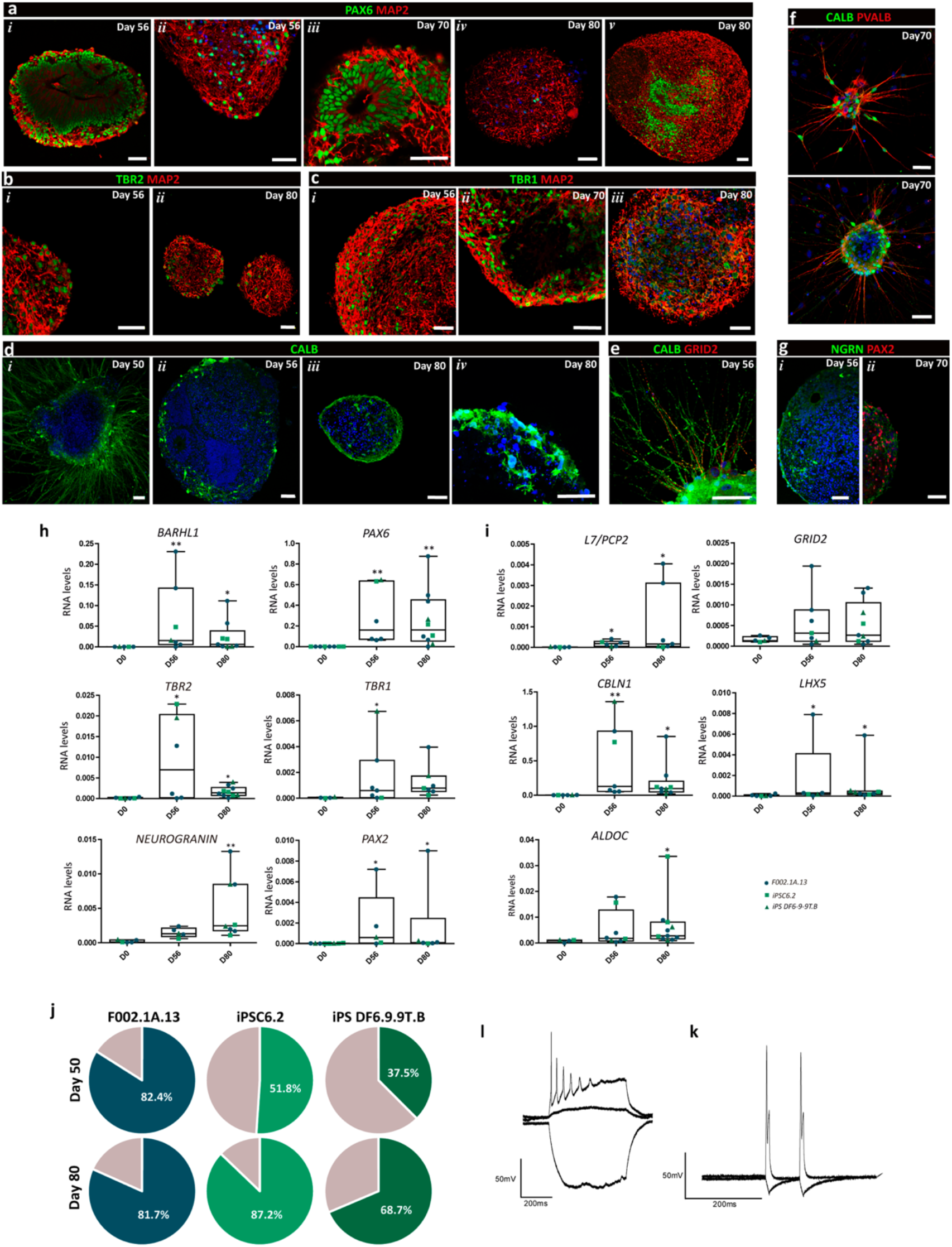
Different types of functional cerebellar neurons in human iPSC-derived aggregates. **a-g,** Immunofluorescence of distinct proteins (as indicated at the bottom of each image) on indicated time-points of differentiation. Scale bars, 50μm. **h-i,** qRT-PCR analysis of aggregates derived from different human iPSC lines at the indicated time-points for markers of different types of cerebellar neurons, including Purkinje cell markers (i). Box-plots diagrams depict mRNA expression levels (2^-ΔCT^) relative to *GAPDH*. Each color and shape-coded dot represents data from one differentiation experiment. One-way ANOVA (Kruskal-Wallis test), *p<0.05, **p<0.01, ***p<0.001; error bars represent SEM. **j,** Percentage of neurons, displaying a histamine/KCl response ratio below 0.8, is shown in colored slices. **l-k,** Whole path-clamp recording on day 80 of differentiation. Representative traces of firing response evoked by a 500 ms current pulse (l) and firing responses to two independent current injections (10 ms) separated by 80 ms (k).

To evaluate the maturation in cerebellar organoids, we performed single-cell calcium imaging. For that, organoids were gently dissociated, and single cells were stimulated with 50 mM KCl and 100 μM histamine. KCl treatment leads to neuron depolarization increasing intracellular calcium concentration, whereas histamine stimulation increases calcium concentration in progenitor cells. Thus, we quantified the percentage of KCl responsive cells, presenting a histamine/KCl ratio below 0.8. Quantification of the percentage of neurons in different human iPSC lines demonstrated that this percentage varied between 37.5% and 82.4% within the organoids by day 50 (**Fig. 3j**). Nevertheless, the organoids from all cell lines presented a gradual and time-dependent neuronal differentiation, achieving 69% to 87% of neurons at the end of the differentiation protocol (**Fig. 3j**). These results were consistent with the presence of cell niches containing KI67^+^, SOX2^+^ and PAX6^+^ neural progenitors during all maturation process (**Supplementary Fig. 2b-g**), retained until day 80 of differentiation. Furthermore, the electrophysiological properties of differentiated neurons were evaluated by patch-clamp recordings after organoid dissociation and replating. The differentiated neurons within organoids on day 80 exhibited fire action potential after a continuous current injection (**Fig. 3l**) and were able to respond to two different current injections separated by 10ms (**Fig. 3k**), demonstrating their capability for depolarizing, repolarizing and recovering. Thus, organoids in this dynamic condition were maintained viable for up to 80 days (**Supplementary Fig. 2h**), containing different types of functional cerebellar neurons.

### Dynamic culturing condition allows more efficient mid-hindbrain commitment and faster cerebellar differentiation

A robust cerebellar commitment and differentiation was previously demonstrated using a static culture system(Silva et al., 2020a). However, in static condition we found that organoids tend to coalesce starting from day 21 of differentiation, forming large macroscopic structures with a dense cell mass (**Supplementary Fig. 3a**), which might resist to diffusion of oxygen, nutrients and morphogens. Differently, in VWBRs, organoids exhibited a pronounced epithelization similar to neural tube structures with luminal space (**Supplementary Fig. 3a**). The analysis of diameter distributions reveled a more homogeneous aggregate size during cerebellar differentiation in dynamic condition when compared with static culture (**Supplementary Fig. 3b**). Likewise, at the end of cerebellar commitment, much larger aggregates were present in static condition, reaching more than 1000 μm in some cases (**Supplementary Fig. 3b**).

To determine the impact of dynamic culturing condition on cerebellar commitment and differentiation, a transcriptomic analysis of organoids derived from static and dynamic differentiations was performed. Principal component analysis (PCA) demonstrated considerable transcriptomic differences between organoids obtained in static and dynamic conditions at different timepoints (**Fig. 4a**). The global gene expression profiles showed a significant clustering of samples by conditions and differentiation stage, in which PC1 represents 54% of variance (**Fig. 4a**), probably related with differentiation progress, followed by 27% of variance captured by PC2 (**Fig. 4a**), suggesting differences between static and dynamic conditions. By day 14, GO analysis of significantly upregulated genes in the dynamic protocol (**Fig. 4b-c**) when compared with static conditions showed enrichment of regionalization (GO:0021871; GO:0021978) and pattern specification processes (GO:0009952, **Fig. 4c**), related with the self-organization and patterning of cells. Interestingly, midbrain development processes were enriched in dynamic conditions, confirmed by higher number of normalized read counts of transcripts of representative genes annotated in the midbrain development (**Supplementary Fig. 3c**). Nevertheless, when we evaluated the top 20 genes that contributed for this difference in the normalized read counts, some of the transcripts that were upregulated in dynamic protocol by day 14 are also annotated and reported to be involved in hindbrain development (4/8 genes, **Supplementary Fig. 3d**), suggesting a possible more efficient mid-hindbrain patterning in the dynamic protocol by day 14. In addition, the analysis of transcripts annotated in cerebellar development showed a higher expression in dynamic conditions by day 14 and achieved similar levels at day 35 (**Fig. 4d**), suggesting a faster cerebellar differentiation in the dynamic protocol when compared with static. By qRT-PCR analysis, the mRNA levels of transcripts critical for cerebellar development, including *OTX2*, *EN2*, *PAX2*, *PAX6* and *KIRREL2*, and those expressed in cerebellar progenitors, such as *ATOH1, BARHL1, OLIG2* and *CORL2*, were found up-regulated in dynamic conditions (**Fig. 4e**), supporting an accelerated cerebellar commitment and further differentiation. On the other hand, by day 35 no significant differences were observed in the expression of mRNA levels for *ATOH1, BARLH1, OLIG2* and *CORL2* by qRT-PCR between different culture conditions (**Supplementary Fig. 3e**), as expected and demonstrated in **Fig. 4d**.

**Figure 4.**
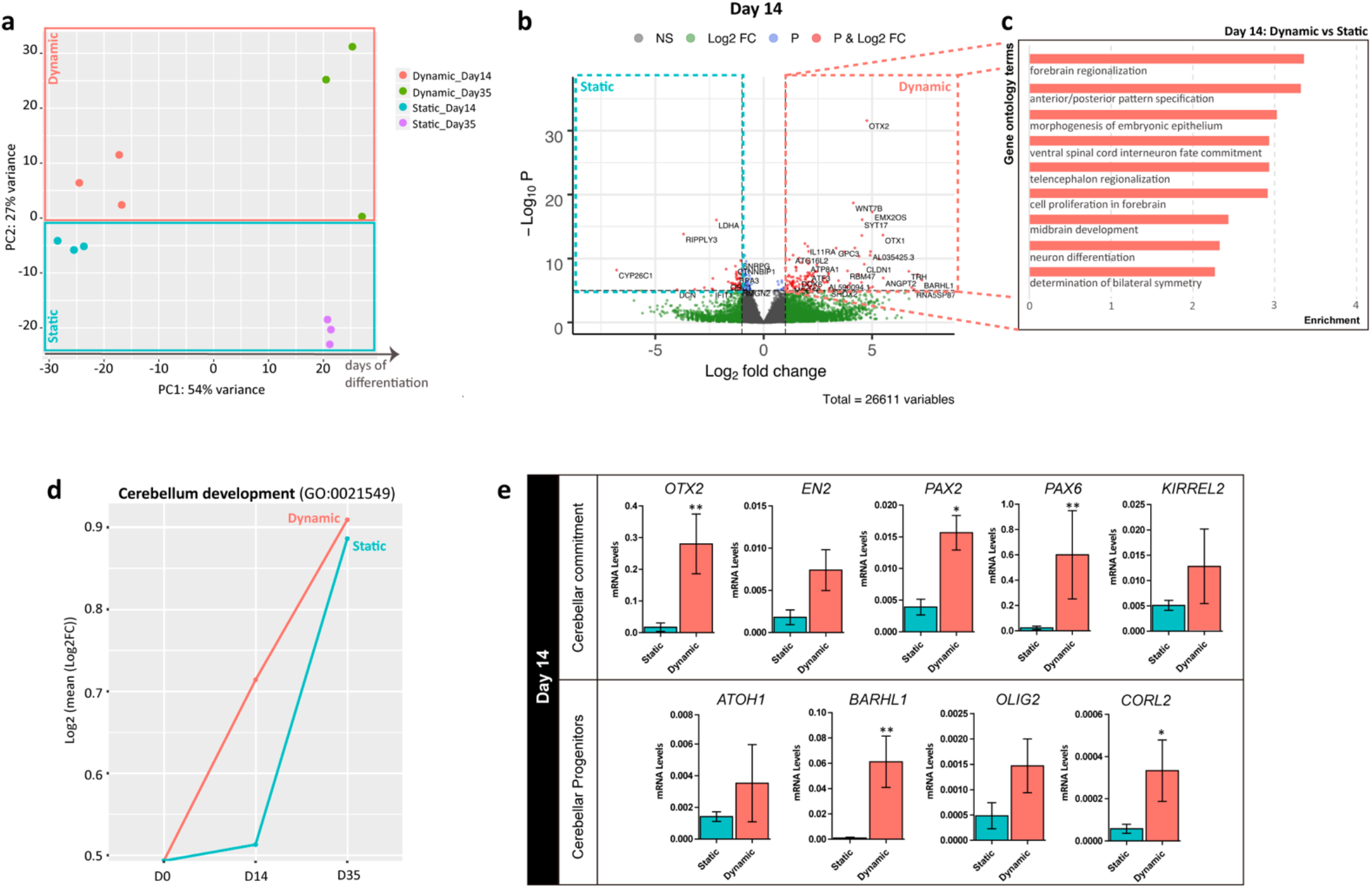
Transcriptomic profiles reveal a more efficient midbrain commitment and accelerated cerebellar differentiation of human iPSCs in the dynamic condition. **a,** PCA based on whole transcriptome data of aggregates cultured in static or dynamic condition at days 14 and 35 of cerebellar differentiation (n=3 for each condition). **b,** Volcano plot of genes identified after cerebellar differentiation of human iPSCs on day 14, comparing static and dynamic conditions. Significantly enriched genes in each condition are labeled. **c,** Top gene ontology (GO) biological processes terms identified for the differentially up-regulated genes (Log2 FC > 2 and adjusted p-value < 0.05) of dynamic versus static condition at day 14 of differentiation. **d,** Line plot summarizing the expression of representative genes annotated in cerebellar development (GO:0021549) relative to day 0 along the differentiation protocol. N=111 genes **e,** qRT-PCR analyses of aggregates derived from F002.1A.13 iPSC line at day 14 of differentiation. Diagrams depict mRNA expression levels (2^-ΔCT^) relative to *GAPDH* based on five independent differentiation experiments for each condition. Mann-Whitney test; *p<0.05, **p<0.01; error bars represent SEM.

### Shear forces promote a significant enrichment of extracellular matrix during cerebellar differentiation

Mechano-transduction and hemodynamic forces represent essential regulators of early differentiation events during embryonic development(Culver and Dickinson, 2010). Therefore, hydrodynamic shear induced by fluid flow may promote stem cell differentiation toward a specific germinal layer, depending on its magnitude and duration(Wolfe et al., 2012). To determine the impact of shear stress on cerebellar organoids, the transcriptome adaptations to dynamic conditions during cerebellar differentiation were analyzed. Gene set enrichment analysis (GSEA) of significantly upregulated genes (Log2 FC > 2 and adjusted p-value < 0.05, **Supplementary Fig. 4a**) during cerebellar differentiation revealed enrichment in pattern specification (GO:0009952), regionalization (GO:0021978) and morphogenesis processes (GO:0048562; GO:0048646) in dynamic culture versus the static counterpart (**Fig. 5a**), as well as biological processes involved in axon guidance and neuron migration (**Fig. 5a**). Extracellular matrix (ECM) organization was detected in dynamic conditions by GSEA, supported by higher number of normalized read counts of transcripts of representative genes annotated to the ECM organization in dynamic condition by days 14 and 35 (**Supplementary Fig. 4b**). Analysis of differentially expressed transcripts responsible for ECM organization in dynamic conditions revealed high levels of the proteoglycan *ACAN* and the link protein *TNC* (**Fig. 5b**), both of which are reported to be highly expressed in the neural ECM(Soleman et al., 2013). *SULF1*, which encodes for sulfatase 1 playing an important role in the neurite outgrowth during postnatal cerebellar development(Kalus et al., 2015), was found upregulated in the dynamic protocol. *LAMA2B*, the only laminin subunit identified to be enriched in the dynamic protocol (**Fig. 5b**), has been demonstrated to be needed for proper synapse assembly(Sann et al., 2008). In addition to this laminin subunit, different genes involved in integrin cell surface interactions (HSA-216083) were found enriched in dynamic culture, including *FBN1, ITGA2, ITGA8* and *ITGB5* (**Fig. 5b**). Furthermore, dynamic cultures of cerebellar organoids presented a collagen-rich ECM, with increased expression of *COL4A5, COL6A3* and *COL9A1* transcripts (**Fig. 5b**), involved in NCAM signaling for neurite out-growth (HSA-375165) and axon guidance (HSA-422475) processes. Genes belonging to ECM (GO:0030198; GO:0031012; **Supplementary Fig. 4c**) and extracellular region (GO:0005576) components GO terms were particularly upregulated at day 35 for dynamic conditions (**Supplementary Fig. 4c**). The presence of extracellular components was further evaluated by immunostaining on day 35, revealing the expression of LAMININ (**Fig. 5c, *i***), FIBRONECTIN (**Fig. 5c, *ii* and *iv***) and COLLAGEN I (**Fig. 5c, *iii* and *iv***), which was maintained until the end of differentiation (**Supplementary Fig. 4d**). Furthermore, AGGRECAN (**Fig. 5c, *v***) and VERSICAN (**Fig. 5c, *vi***), which are heparin sulfate proteoglycans highly expressed in neural ECM(Howell and Gottschall, 2012), were also observed by immunostaining on day 35 along with SYNAPSIN (**Fig. 5c, *v* and *vi***), a presynaptic phosphoprotein fundamental for the regulation of synaptic transmission. This demonstrates a possible connection between synaptic processes and some elements of the ECM(De Luca et al., 2020). In addition to ECM organization, cerebellar differentiation in dynamic conditions also appeared to enhance cell adhesion processes when compared with static culturing (**Fig. 5a**). The differentially expressed genes annotated in cell adhesion process that were found upregulated in dynamic culture include *ALCAM, BCL2, EPHA7, FLRT2, LAMB2, LRRN2, PLXNA4, RELN, TNC* and *UNC5D* transcripts (**Fig. 5d**), which are also involved in axon (GO:0061564) and neuron projection development (GO:0031175), supporting an efficient neuronal differentiation. Interestingly, in addition to neuronal transcripts, cell adhesion genes involved in circulatory system development (GO:0072359), including *ACVR1, ADAMTS9, ANGPT2, BMP2, BMP4, COL1A1, DSP, EPHA2, FBN1, FLRT2, FOXJ1, PPARA, CD36* (**Fig. 5d**), and regulation of vasculature development (GO:1901342), like *ADAMTS9, ANGPT2, BMP4, EPHA2, WNT4* (**Fig. 5d**), were found differentially expressed in organoids derived from dynamic culture when compared with static. Thus, fluid flow established by agitation in the dynamic protocol may activate additional transcriptional regulation of genes involved in angiogenesis(Wolfe and Ahsan, 2013). Some significantly downregulated genes (Log2 FC < −2 and adjusted p-value < 0.05) in dynamic culture versus the static counterpart during cerebellar differentiation were detected, however none biological processes associated with these genes were strongly enriched.

**Figure 5.**
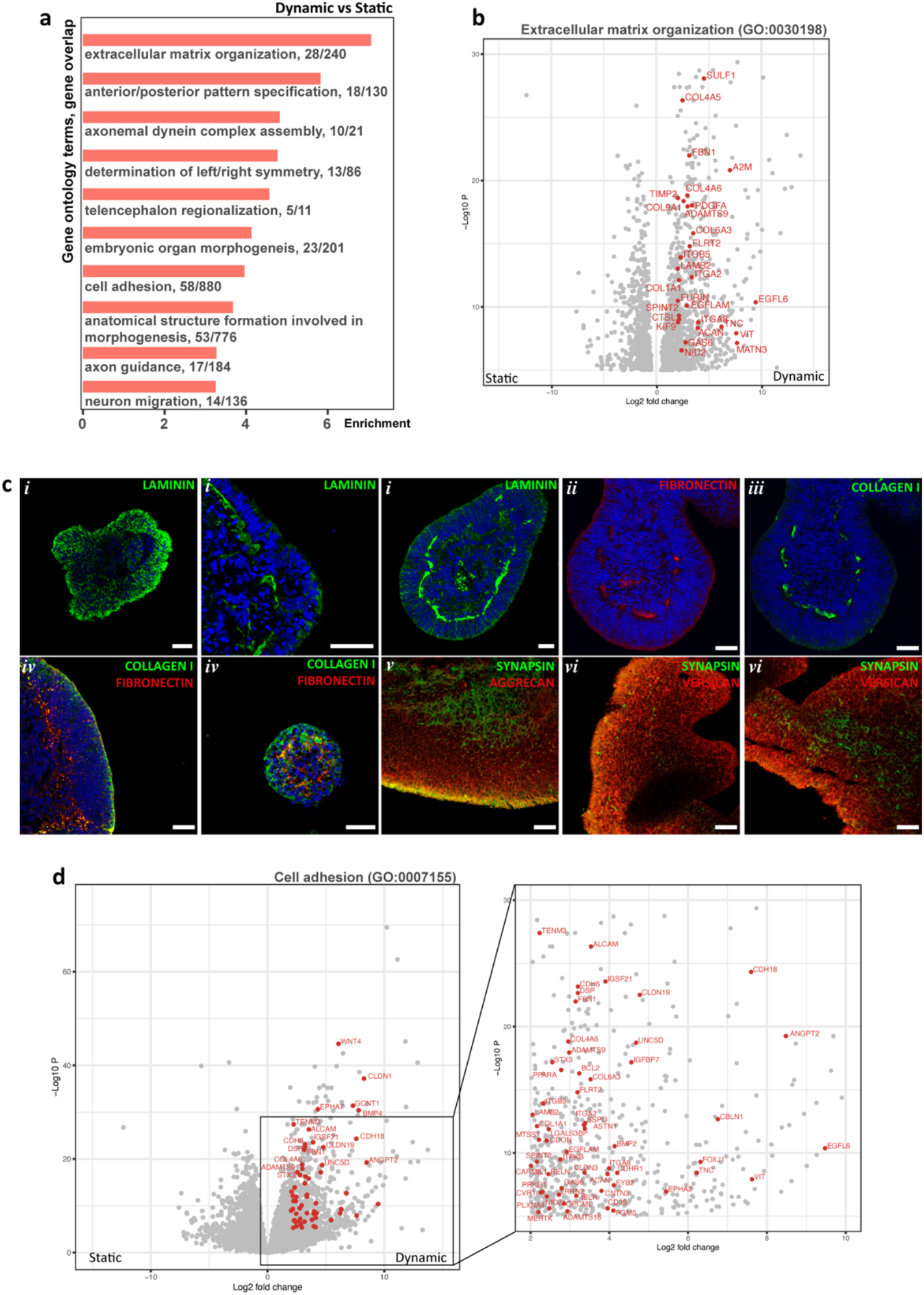
Global comparison of organoids transcriptome reveals a significant remodeling of extracellular matrix (ECM) in dynamic conditions when compared with static protocol. **a,** Top gene ontology (GO) biological processes terms identified for the differentially up-regulated genes (Log2 FC > 2 and adjusted p-value < 0.05) of dynamic versus static conditions in cerebellar differentiation. **b,** Volcano Plot of differentially expressed genes between dynamic and static conditions showing significantly up-regulated transcripts in dynamic conditions (Log2 FC > 2 and adjusted p-value < 0.05) annotated in the ECM organization process (GO:0030198). **c,** Immunostaining analysis on days 35 of differentiation in dynamic conditions for the indicated ECM markers. Scale bars, 50μm. **d,** Volcano Plot of differentially expressed genes between dynamic and static conditions showing significantly up-regulated genes in dynamic conditions (Log2 FC > 2 and adjusted p-value < 0.05) annotated in the cell adhesion process (GO:0007155).

### Transcriptional modulation of genes involved in angiogenic processes by dynamic condition

Neurogenesis and angiogenesis are two different processes that seem to be tightly coupled during embryonic neural development. Nascent blood vessels were reported to actively contact dividing neural stem cells and have a function in their behavior, impacting proper brain development (Di Marco et al., 2020). To further understand how culturing cerebellar organoids in dynamic condition impacts on angiogenic transcriptional profiles, we analyzed the number of normalized read counts of representative genes annotated in the sprouting angiogenesis process (GO:0002040). An increased expression of these transcripts was detected in dynamic condition by day 35 (**Fig. 6a**). Among the top 20 genes that contributed the most to the observed differences, *BMP4, ESM1*, *EPHA2* and *ADAMST9* transcripts were detected at high levels in dynamic cultures by day 35 (**Supplementary Fig. 5a**). While *ESM1* and *EPHA2* are essential regulators of angiogenesis, modulating endothelial cell behavior and migration (Brantley-Sieders et al., 2004; Rocha et al., 2014), *ADAMST9* is expressed by microvascular endothelial cells (Koo et al., 2010), being abundantly present in the central nervous system (CNS, Gottschall and Howell, 2015). The expression of these key genes was confirmed by qRT-PCR, showing a significantly higher mRNA levels in dynamic condition in comparison with static culture at day 35 of differentiation (**Fig. 6b**), which were maintained at high levels until day 80 in 3D cerebellar organoids when compared with day 0 (**Supplementary Fig. 5b**). The proteins encoded by these transcripts have been reported to act by binding directly to the ECM (Rocha et al., 2014). Thus, we next assessed the transcriptional modulation of ECM organization and sprouting angiogenesis processes across cerebellar differentiation using static and dynamic protocols. Interestingly, the expression variation of transcripts annotated in these two processes was very similar, with an increased expression from day 0 to 14 in both conditions (**Fig. 6c**). Thereafter, expression dropped in static conditions and increased until day 35 in the dynamic protocol (**Fig. 6c**). Immunostaining also confirmed the presence of CD31^+^ cells among LAMININ^+^ ECM (**Fig. 6d**), suggesting a possible interaction between ECM enrichment and angiogenesis onset. Additionally, this dynamic condition was significantly enriched for collagen trimer and blood vessel development-specific gene signature (**Supplementary Fig. 5c**), confirmed by staining for CD34 and COLLAGEN I (**Supplementary Fig. 5d**). Sprouting angiogenesis is reported to be usually initiated by hypoxia and afterwards the maturation and stabilization of capillaries requires ECM deposition, as well as shear stress and other mechanical signals (Chien, 2007). Therefore, we evaluated the enrichment of transcripts involved in response to mechanical stimulus (GO:0009612) and cellular response to hypoxia (GO:0071456). In contrast to cellular response to hypoxia, where no substantial differences were identified between static and dynamic condition (**Fig. 6e**), genes annotated in response to mechanical stimulus were essentially enriched in the dynamic protocol (**Fig. 6e**). Interestingly, HIF1*α* transcriptional activation was found downregulated in the dynamic protocol in comparison with static conditions, based on top 100 genes (**Supplementary Fig. 5e**), ruling out the contribution of hypoxia to activate the angiogenic process in the dynamic system. Based on these results, the onset of angiogenesis in the dynamic condition might be triggered by two different signaling pathways, either by ECM or mechanical stimulus (**Fig. 6f**). In the CNS, angiogenesis seems to be promoted by ECM signaling supported by NRP1 expression, which interacts with ABL1 and CDC42, triggering the angiogenic process in a VEGF-independent manner (Fantin et al., 2015; Raimondi et al., 2014) (**Fig. 6f**). On the other hand, shear stress was reported to stimulate the expression of VEGFR2 mediated by induction of KLF2 (dela Paz et al., 2012; Renz et al., 2015) (**Fig. 6f**). Thus, we quantified the mRNA levels of these two candidates, *NRP1* and *KLF2*, as well as *VEGFA*, to evaluate the contribution of ECM and shear stress to the increased angiogenic pathway expression observed in dynamic cerebellar organoid cultures. *NRP1* and *KLF2* transcripts show a marked increase throughout the dynamic differentiation protocol, achieving significantly higher levels in dynamic condition in comparison with static by day 35 (**Fig. 6g**). Despite significant differences in *KLF2* mRNA levels between static and dynamic protocols, no differences in the expression of *VEGF2A* mRNA was observed by day 35 (**Fig. 6g**). From day 35, while *NPR1* mRNA levels were maintained constant until the end of the cerebellar differentiation (**Fig. 6h**), *KLF2* transcript expression continued to increase significantly until day 80 (**Fig. 6h**), which was accompanied by a slight but statistically significant increase of *VEGFA* levels from day 14 until day 80 of differentiation (**Fig. 6h**). Therefore, a possible combination of ECM and mechano-transduction pathways may contribute for the onset of the angiogenic fate, which leads to a significant expression of CD31 and CD34 until day 80 of cerebellar differentiation (**Supplementary Fig. 5f-g**). Human protein-protein interactome network for genes responsible for the modulation of angiogenic processes observed during differentiation confirmed that transcripts annotated in integrin-mediated signaling and response to stress play an active role in the regulation of angiogenesis, however, they also have an important function in neuronal processes such as, for example, neuron projection morphogenesis (**Fig. 6i**). Interaction between different gene clusters demonstrated that a common activation of neuronal and angiogenic fate may be achieved throughout the cerebellar differentiation protocol, and we hypothesize that this may be important for the higher degree of neuronal differentiation observed in dynamic condition.

**Figure 6.**
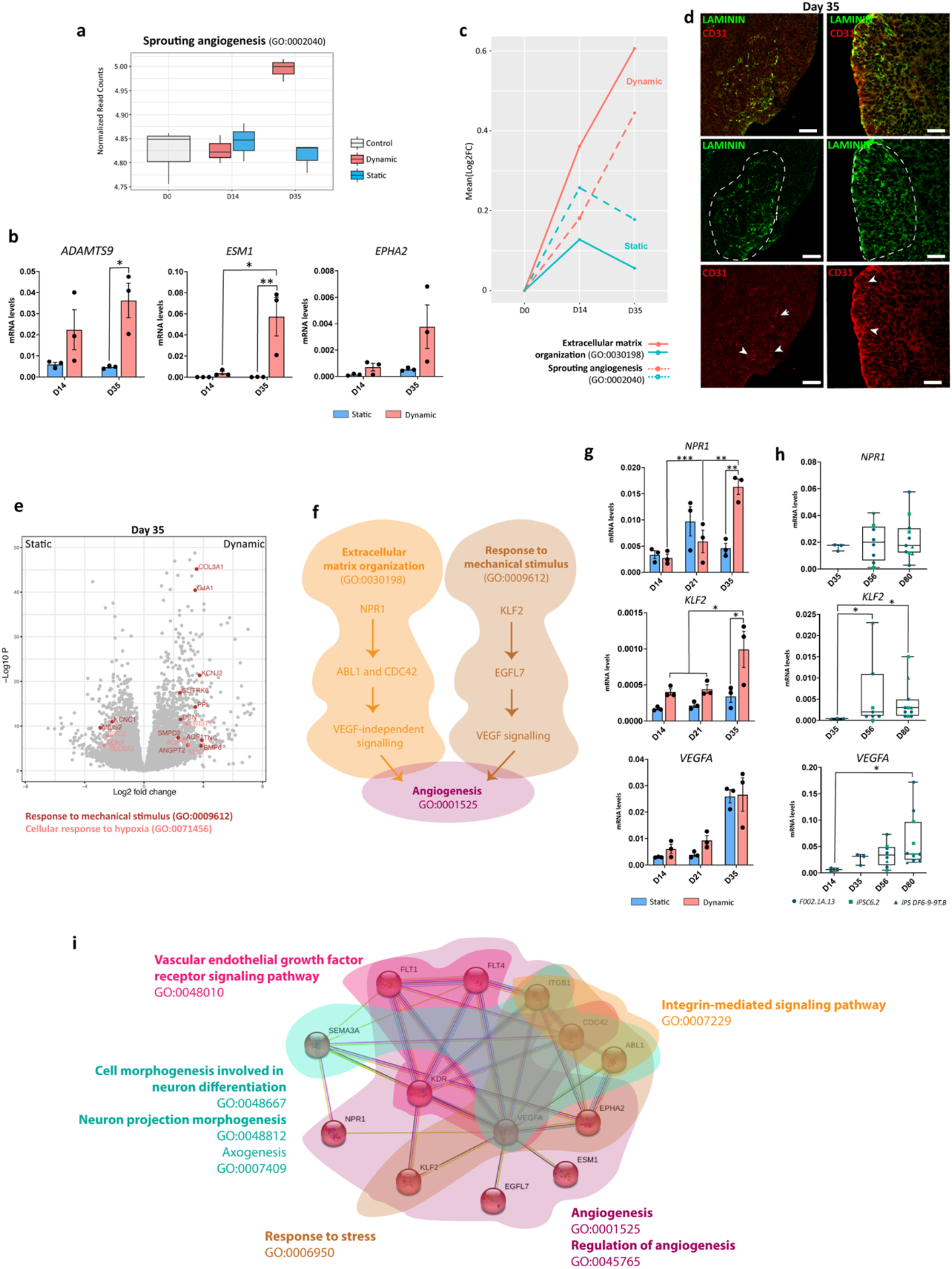
Dynamic condition modulates the expression levels of genes that favor the angiogenesis onset. **a,** Box-plot representing RNA-seq normalized read counts of representative genes annotated in the sprouting angiogenesis process (GO:0002040, n=125 genes) for control (day 0), dynamic and static conditions at indicated time-points of differentiation. **b,** qRT-PCR analyses of organoids derived from F002.1A. 13 iPSC line for genes involved in angiogenesis process. Diagrams depict mRNA expression levels (2^-ΔCT^) relative to GAPDH based on three independent differentiation experiments for each condition. Two-way ANOVA (Tukey’s multiple comparisons test), *p<0.05, **p<0.01; error bars represent SEM. **c,** Line plot representing the expression of representative transcripts annotated in ECM organization (GO:0030198, n=240 genes) and sprouting angiogenesis (GO:0002040 n=125 genes) processes for static and dynamic conditions relative to day 0. **d,** Immunofluorescence for LAMININ and CD31 on day 35 of dynamic differentiation. Scale bars, 50μm. **e,** Volcano Plot of differentially expressed genes between dynamic and static conditions showing significantly down- and up-regulated transcripts in dynamic conditions (|Log2 FC| > 2 and adjusted p-value < 0.05) related with response to mechanical stimulus (GO:0009612, n=180 genes) and cellular response to hypoxia (GO:0071456, n=180 genes) processes. **f,** Schemes illustrating the proposed networks that may be involved in the modulation of angiogenic process by either independent or dependent-VEGF signaling pathway. **g,** qRT-PCR analyses of organoids derived from F002.1A.13 iPSC line. Diagrams depict mRNA expression levels (2^-ΔCT^) relative to GAPDH based on three independent differentiation experiments for each condition. Two-way ANOVA (Tukey’s multiple comparisons test), *p<0.05, **p<0.01; error bars represent SEM. **h,** qRT-PCR analysis of organoids derived from different human iPSC lines at the indicated time-points. Box-plots diagrams depict mRNA expression levels (2^-ΔCT^) relative to GAPDH. Each color and shape-coded dot represents data from one differentiation experiment. One-way ANOVA (Kruskal-Wallis test), *p<0.05; error bars represent SEM. **i,** Protein-protein interaction network prediction based on protein encoding genes involved in angiogenic process. Interaction networks were obtained from STRING database, in which nodes were clustered using Markov Cluster Algorithm (MLC) with an inflation parameter = 3. Biological processes (FDR < 0.05) annotating the analyzed proteins are highlighted.

## Discussion

We reported a dynamic culture system able to generate human iPSC-derived cerebellar organoids, which matured into functional cerebellar neurons, using the novel VWBRs. The large-scale production of cerebellar organoids represents an important advance for automated high-throughput drug screening, as well as in regenerative medicine applications for neurodegenerative diseases that affect the cerebellum. Previous studies have mainly reported the scalable differentiation of PSC into neural progenitors and functional neurons (Bardy et al., 2013; Miranda et al., 2016; Rigamonti et al., 2016), lacking the recapitulation of structural cell organization seen during human embryonic development, as well as the ability to maintain these 3D structures containing functional neurons for long periods of time.

### Dynamic condition allows a reproducible and large-scale generation of organoids

By using the VWBRs, we applied a novel mixing mechanism generated by a large vertical wheel that rotates around the horizontal axis, and allows gentle and uniform fluid mixing (Croughan et al., 2016). Therefore, we ensured that uniform exposure of neural organoids to signaling molecules was reached. Moreover, the operation time and complexity, as well as the risk of contaminations, is reduced by using these single-use vessels, which facilitate the adoption of GMP conditions (Croughan et al., 2016). With this dynamic culture system, we were able to address several critical issues. First, it allows to easily generate a high number of organoids, which is important for high-throughput applications, attaining around 350 ± 52 (mean ± SEM) aggregates/mL in the first 24 hours after iPSC seeding at 250 000 cells/mL (**Fig. 1d** and **Supplementary Fig. 1b**). Beyond that, it is important to ensure that the seeding process leads to maximum cell survival and homogeneous aggregate production, since the aggregate size has a critical role in inducing differentiation towards a specific cell lineage (Bauwens et al., 2008). Indeed, a high percentage of cell survival was observed after the single cell seeding, once 54.1 ± 9.3%, 80.7 ± 2.4% and 88.2 ± 11.8% of live cells were able to aggregate using three independent human iPSC lines (**Supplementary Fig. 1b**). Moreover, aggregates formed in VWBRs were uniform in size and shape, achieving the optimal diameter (Miranda et al., 2015; Miranda et al., 2016; Silva et al., 2020a) to initiate neural induction at 48 hours after cell seeding (**Fig. 1b-c**). In comparison to static condition, organoids retained a more homogeneous diameter in VWBRs during the differentiation protocol (**Supplementary Fig. 3a-b**), which is useful to reduce variability between multiple experiments.

### Cerebellar organoids differentiated in dynamic conditions recapitulate human cerebellar structure

Using size-controlled human iPSC-derived aggregates and a chemically defined medium, we were able to recapitulate sequential steps of human cerebellar development in a continuous differentiation process, starting with an efficient cerebellar commitment (**Fig. 2**), and further differentiation into different types of cerebellar neurons (**Fig. 3**). By immunostaining and gene expression analysis, we successfully and readily identified distinct types of cerebellar Glutamatergic and GABAergic neurons (**Supplementary Fig. 2a**), as seen in layers of human cerebellar cortex and in cerebellar nuclei. Specifically, the following cell types can be produced in our dynamic culture system: Granule neurons (Pax6^+^/MAP2^+^), Unipolar Brush cells (TBR2^+^/MAP2^+^), DCN projection neurons (TBR1^+^/MAP2^+^), Purkinje cells (Calbindin^+^/GRID2^+^), non-Golgi GABAergic interneurons (Calbindin^−^ /Parvalbumin^+^) and Golgi cells (Neurogranin^+^/PAX2^+^) (**Fig. 3a-g**). Furthermore, calcium imaging and electrophysiological evaluation indicate that cerebellar precursors have achieved an efficient maturation in our dynamic cultures (**Fig. 3j-k**). Interestingly, in addition to the functional establishment of neuronal network connectivity, several pools of neural progenitors were maintained until day 80 in culture during the maturation process. The maintenance of a neural progenitor niche during CNS development is an important biological process achieved by the microenvironmental cues, as well as cell-cell interactions, which are capable to balance stem cell quiescence with proliferation and to direct neurogenesis versus gliogenesis (Conover and Notti, 2008). Also, during cerebellar development, different niches of cerebellar progenitor cells are observed, the first in the ventricular zone and the second in the external granule layer (ten Donkelaar et al., 2003; Wingate, 2001). SOX2^+^ progenitor niche is present in the ventricular zone, where this protein is highly expressed from gestation weeks 20 to 24, being downregulated in the developing human cerebellum, with undetectable expression by week 38 (Pibiri et al., 2016). On the other hand, an external granular layer expressing PAX6 is observed from weeks 20 to 38 of gestation (Pibiri et al., 2016). Therefore, our 3D culture system seems to recreate the *in vivo* microenvironment observed between gestation weeks 20 and 24, supported by the presence of SOX2^+^ and PAX6^+^ progenitor niches (**Supplementary Fig. 2b-f**).

### Faster and efficient cerebellar differentiation is promoted by dynamic culture

The VWBRs allowed us to establish a scalable and efficient system for human iPSC cerebellar commitment, with a homogeneous culture environment inside the vessel and complete suspension of cell aggregates. However, despite the particularly gentle mixing mechanism of the VWBRs, in bioreactor cultures cells are exposed to hydrodynamic shear stress inherent to the suspension culture.

As it was already reported that shear stress is an important regulator of germinal specification (Kumar et al., 2017; Wolfe et al., 2012), we analyzed the transcriptional changes occurring during cerebellar differentiation using dynamic conditions and compared with the static control. Evaluating the transcriptomic profiles in the initial steps of cerebellar induction, we detected a more efficient mid-hindbrain commitment, as well as a faster cerebellar differentiation in dynamic conditions (**Fig. 4** and **Supplementary Fig. 3**). The initial mid-hindbrain patterning and further cerebellar induction was achieved by using moderate caudalizing factors, like FGF2 and insulin, which were reported to induce the expression of FGF8 and EN2, crucial transcription factors involved in the isthmic organizer (Chi et al., 2003; Martinez et al., 1999) and mid-hindbrain boundary maintenance (Joyner, 1996). Yoshiki Sasai proposed that this combination can efficiently promote the self-formation of isthmic organizer tissue (OTX2^+^/EN2^+^), representing a small area within the organoid (Sasai, 2013). Afterwards, the self-production of signals from this organizer center, like FGF8, stimulate the generation of cerebellar territory (Sasai, 2013). In our dynamic culture, design features of VWBRs (Croughan et al., 2016) provide efficient fluid mixing and enhanced mass transfer within the organoids, promoting a uniform diffusion of signaling molecules, either exogenously provided morphogens or endogenous signals emanating from neighboring cells. A possible explanation for enhanced cerebellar tissue patterning in dynamic conditions may be that iPSC-derived aggregates were exposed to an efficient and uniform diffusion of FGF2 signaling that could result in a larger area of isthmic organizer tissue (OTX2^+^/EN2^+^). On the other hand, the maintenance of this organizer tissue is also dependent on the self-production of FGF and WNT signaling, as well as their suppression by inhibitors, in a reaction-diffusion model (Kondo and Miura, 2010; Turing, 1952). Thus, dynamic culture conditions, with more efficient fluid mixing, can lead to uniform exposure to signaling cues and enhance the feedback loops that operate in this self-organized system.

### Dynamic cerebellar differentiation induces ECM enrichment

In our time-course data, comparing dynamic and static conditions, we detected that significant ECM enrichment was promoted by using VWBRs for cerebellar differentiation, diverging from the transcriptomic profiles of static condition (**Fig. 5**). Indeed, ECM components are not only highly expressed within neural tissues, but also have a great impact in some aspects of neural development (Jain et al., 2020; Long and Huttner, 2019). With this significant enrichment of ECM in our dynamic cultures, we created a system that avoids the use of exogenous ECM, for instance encapsulation with Matrigel, that is commonly used to support the maintenance of brain organoids in spinner flaks (Lancaster et al., 2012; Qian et al., 2018), being a source of a significant heterogeneity between organoids (Nayler et al., 2020). In our dynamic system, self-derived ECM was enriched in proteoglycans, including aggrecan and versican, as well as a link protein (tenascin C), all of which are highly expressed in neural ECM. Besides that, transcriptomic data analysis of dynamic cultures shows a significant expression of *SULF1*, which supports neurite outgrowth during cerebellar development (Kalus et al., 2015). Other ECM components involved in synaptic processes, neurite outgrowth and axon guidance were highly enriched in dynamic cultures when compared with static, sustaining an efficient neuronal differentiation. In fact, significant changes in ECM during 3D differentiation when compared with 2D were already reported, using perfusion stirred-tank bioreactors for iPSC-derived neural progenitor cells (Simão et al., 2018), confirming that 3D culture associated with dynamic conditions may better mimic protein composition as well as the neural microenvironment.

### Angiogenic process is activated by dynamic culture in cerebellar organoids

In addition to ECM organization, the analysis of global transcriptional modulation across the differentiation protocol also exposed a significant enrichment of transcripts involved in cell adhesion processes (**Fig. 5**). As expected, most of them were involved in neuronal processes, including axon and neuron projection development. Unexpectedly, not only cell adhesion transcripts involved in neuronal development were enriched in dynamic conditions, but also regulators of circulatory system and vascular development. In fact, previous studies have already shown that shear stress employed early in ESC differentiation favored hematopoietic and endothelial fates (Wolfe and Ahsan, 2013). Besides that, the application of shear stress to endothelial cells stimulates the expression of VEGFR2, resulting in increased endothelial cell survival (dela Paz et al., 2012). This may explain the presence of transcripts associated with angiogenic fate upon mechanical stimulation. ECM plays a crucial role in the regulation of the angiogenic process, and the proangiogenic effect of some ECM molecules and fragments (Neve et al., 2014; Stupack and Cheresh, 2002), including Collagen and laminin subtypes, fibronectin and tenascin-C, have been shown. Interestingly, our transcriptomic analysis revealed that the increased expression of ECM components during cerebellar differentiation was complemented by higher expression of transcripts involved in sprouting angiogenic processes (**Fig. 6c**), which was confirmed by co-localization of ECM components and CD31^+^/CD34^+^ endothelial cells (**Fig. 6d**). To investigate a possible origin of this angiogenic process in our dynamic cultures, we analyzed three different mechanisms that could activate angiogenesis: ECM signaling, response to mechanical stimulus and cellular response to hypoxia (Pugh and Ratcliffe, 2003). Hypoxia-based is unlikely, since no differences were found in the expression of transcripts involved in the cellular response to hypoxia between static and dynamic differentiations (**Fig. 6e**). Moreover, HIF1*α* pathway transcripts were included in the top 100 genes downregulated in dynamic conditions comparatively to the static ones **(Supplementary Fig. 5e**). Therefore, two different mechanisms may be responsible for promoting the angiogenic process in our dynamic cultures: one related with ECM organization and the other with cellular response to mechanical stimulus (**Fig. 6f**). In the CNS, angiogenesis is promoted by VEGF-independent signaling, activated by the expression of NPR1 (Tata et al., 2015). NPR1, neuropilin-1, is a non-catalytic transmembrane protein, which was originally identified as an adhesion molecule in the CNS (Takagi et al., 1995) and seems to have an essential role in the vascularization of the hindbrain (Gerhardt et al., 2004). NPR1 was reported to enforce ECM signaling (Raimondi, 2014), in which the extracellular protein domain complexes with integrins, promotes the recruitment of ABL1 and enables CDC42-actin rearrangement independently of VEGF stimulation (**Fig. 6f**), promoting angiogenesis (Fantin et al., 2015; Raimondi et al., 2014). On the other hand, mechano-transduction signaling is also responsible for mediating the angiogenic process in a VEGF-dependent manner (**Fig. 6f**). KLF2 is one of many vasoactive endothelial genes that suffer transcriptional modulation during embryonic development after the establishment of fluid flow (Lee et al., 2006). Moreover, it was demonstrated that shear-stress-induced VEGF expression is mediated by KLF2 expression (dela Paz et al., 2012) and its expression mirrors the rise of fluid shear stress across development *in vivo* (Lee et al., 2006). Overexpression of KFL2 was reported to promote angiogenesis through the upregulation of EGFL7 (Renz et al., 2015), which was found expressed in the vasculature (Parker et al., 2004). Thus, we analyzed the expression of these two different angiogenic inductors, NFR1 and KLF2, to unveil the contribution of ECM remodeling and mechano-transduction signaling, in addition to VEGFA, to confirm whether angiogenesis was promoted by either an independent or dependent-VEGF signaling. Our data show that a possible combination of these two different signaling pathways could contribute for the angiogenesis enhancement observed in our dynamic culture. Significant differences of *NFR1* and *KLF2* transcript expression were found between static and dynamic conditions, with higher mRNA levels presented in dynamic conditions until day 35 (**Fig. 6g**), along with a significant expression of *VEGFA* in dynamic culture-derived organoids from day 14 to 80 of differentiation (**Fig. 6h**). We hypothesize that this combined effect observed in dynamic conditions may be important for the higher degree of neuronal maturation present in our cerebellar organoids (**Fig. 6i**).

## Conclusion

In conclusion, we have reported here the successful and efficient scalable generation of cerebellar organoids, demonstrated by the transcriptomic profile during differentiation. The transcriptomic signatures under dynamic culture conditions revealed significant ECM organization that can better mimic the neural microenvironment and maintain the organoids healthy for at least 80 days of differentiation. One current limitation for the generation of brain organoids has been the recapitulation of a complex microenvironment that involves interaction between different cell types, including vascular cells, neurons, astrocytes, oligodendrocytes and microglia. Here, we demonstrated that dynamic culture can activate signaling pathways important to induce the angiogenic process during neural commitment, introducing significant cues for the recapitulation of a more complex tissue, comprising different cell types. Additionally, this dynamic system allows increased reproducibility between experiments and the possibility of further upscaling the production of cerebellar organoids. Bioreactor technology also has the potential to allow further improvements to the system here described, namely automated monitoring and control of the culture environment (eg. pH or dissolved oxygen concentration) or the use of alternative culture medium feeding strategies such as perfusion (Kropp et al., 2016). We expect that the methodologies developed here will widen the applicability of cerebellar organoids in high-throughput screening, including drug and toxicological testing, as well as in the study of important aspects of pathological pathways involved in cerebellar dysfunction.

## Acknowledgments

This work was supported by Fundação para a Ciência e a Tecnologia (FCT), Portugal (UIDB/04565/2020 through Programa Operacional Regional de Lisboa 2020, Project N. 007317, PD/BD/105773/2014 to T.P.S. and SFRH/BD/147906/2019 to R.S.L.), projects co-funded by FEDER (POR Lisboa 2020—Programa Operacional Regional de Lisboa PORTUGAL 2020) and FCT through grant PAC-PRECISE LISBOA-01-0145-FEDER-016394 and CEREBEX Generation of Cerebellar Organoids for Ataxia Research grant LISBOA-01-0145-FEDER-029298.

## Supplementary data

**Supplementary Figure 1.**
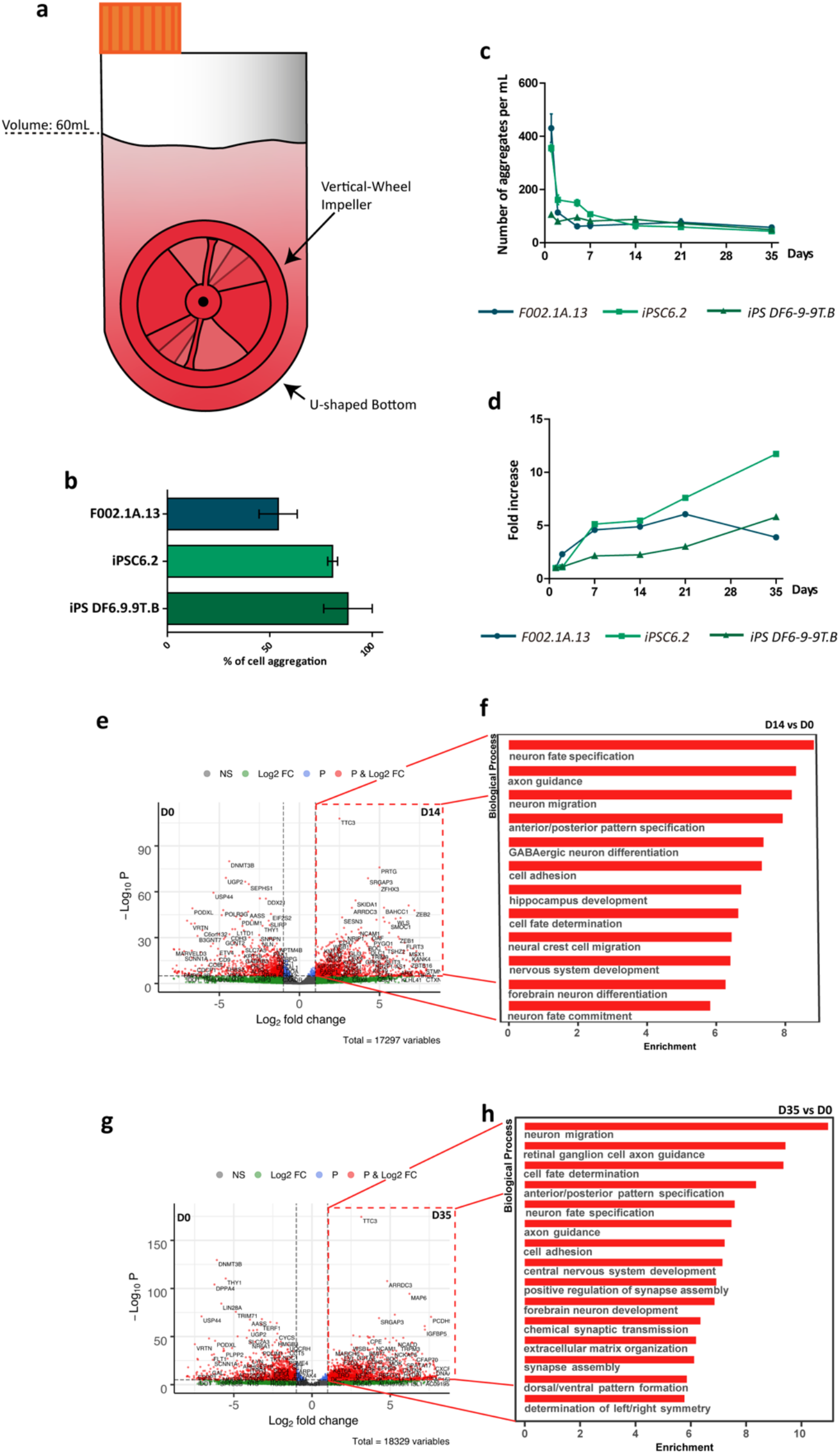
Characterization of human iPSC-derived cerebellar organoids. **a,** Design of vertical-wheel bioreactor. **b,** Percentage of cell aggregation at 24 hours after single-cell inoculation for different iPSC lines. Each bar includes data from 4, 3, and 2 independent differentiation experiments using F002.1A.13, iPSC6.2 and iPS DF6.9.9T.B cell line, respectively. Data are represented as mean ± SEM. **c,** Number of iPSC-derived aggregates from day 1 to day 35 of cerebellar differentiation for each human iPSC line. **d,** Total volume of biomass relative to day 1 using different human iPSC lines. **e,** Volcano plot of genes identified after cerebellar differentiation of human iPSC using VWBRs on day 14 and compared with day 0. Significantly enriched genes after 14 days of differentiation are labeled. **f,** Top gene ontology (GO) biological processes terms identified for the differentially up-regulated genes (Log2 FC > 2 and adjusted p-value < 0.05) of day 14 versus day 0 of differentiation. **g,** Volcano plot of genes identified after cerebellar differentiation of human iPSC using VWBRs on day 35 and compared with day 0. Significantly enriched genes after 35 days of differentiation are labeled. **h,** Top gene ontology (GO) biological processes terms identified for the differentially up-regulated genes (Log2 FC > 2 and adjusted p-value < 0.05) of day 35 versus day 0 of differentiation.

**Supplementary Figure 2.**
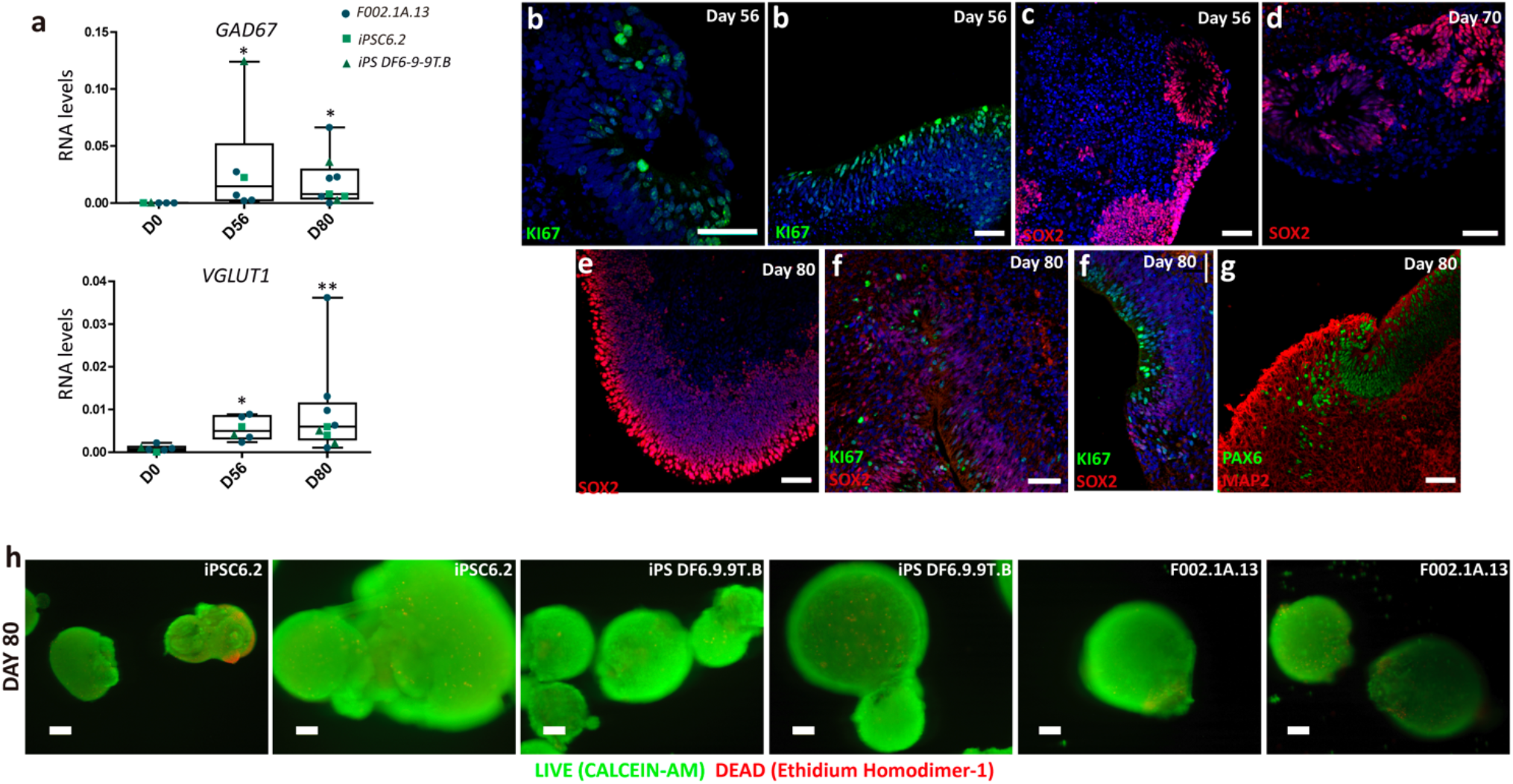
Assessment of maturation and viability in cerebellar organoids. **a,** qRT-PCR analyses of aggregates derived from different iPSC lines at the indicated time-points. Box-plots diagrams depict mRNA expression levels (2^-ΔCT^) relative to GAPDH. Each color and shape-coded dot represents data from one experiment. One-way ANOVA (Kruskal-Wallis test), *p<0.05, **p<0.01; error bars represent SEM. **b-f,** Immunofluorescence for KI67, SOX2, PAX6 and MAP2 on indicated time-points of differentiation. Scale bars, 50μm. **h,** LIVE/DEAD assay for cerebellar organoids derived from distinct human iPSC lines on day 80 of differentiation. Scale bars, 100μm.

**Supplementary Figure 3.**
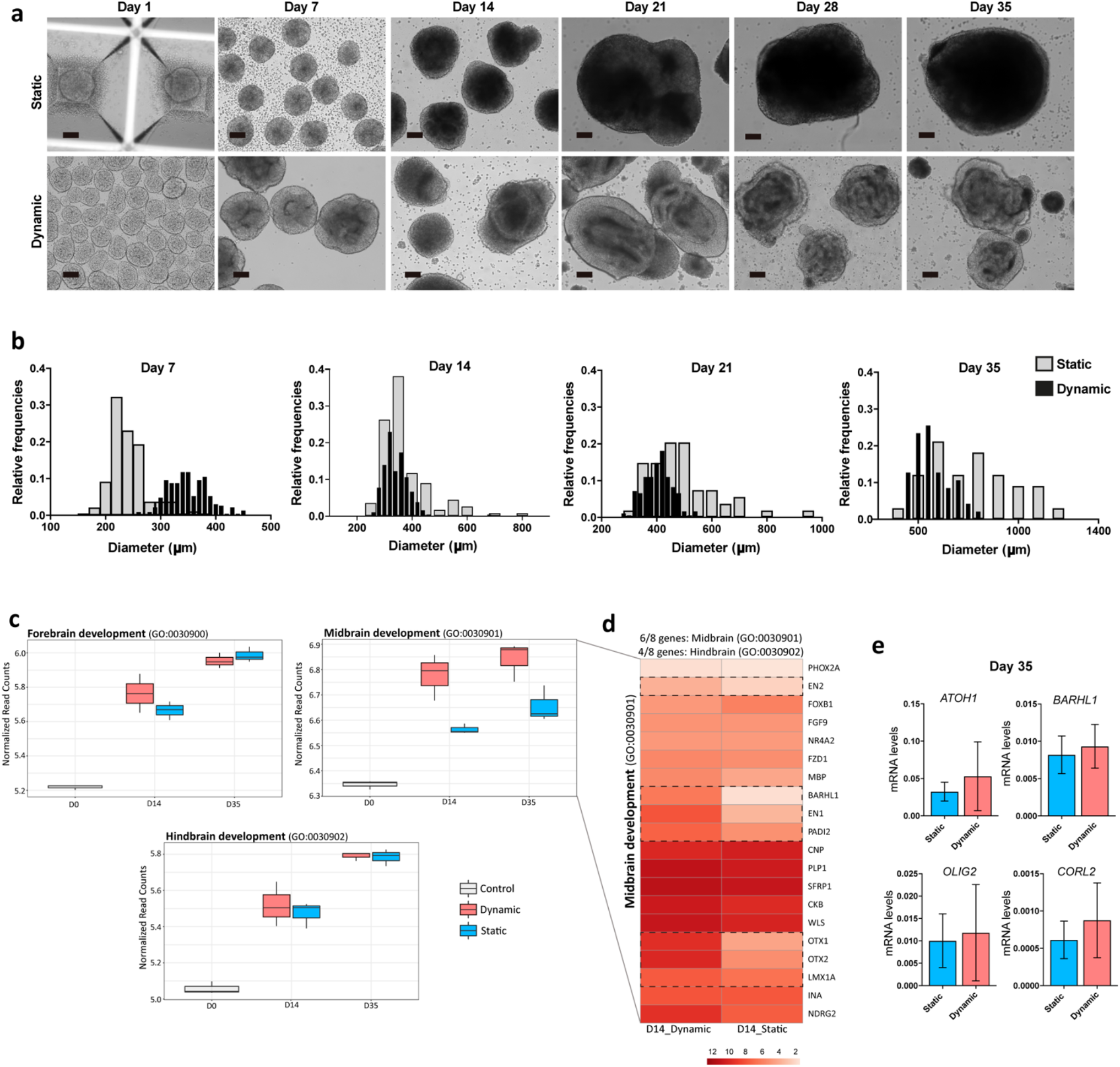
Comparison of dynamic culturing conditions effects on cerebellar differentiation. **a,** Representative brightfield photomicrographs showing human iPSC-derived aggregates during cerebellar differentiation in static and dynamic conditions. Scale bar, 100 μm. **b,** Distribution of aggregate diameters along the differentiation protocol in static and dynamic conditions. **c,** Box-plots summarizing RNA-seq normalized read counts of transcripts annotated in the selected GO terms (forebrain, midbrain and hindbrain development) for control (day 0), dynamic and static conditions at indicated time-points of differentiation. **d,** Heat map highlighting the top 20 differentially expressed genes related with midbrain development (GO:0030901) for dynamic and static conditions at day 14 of differentiation. **e,** qRT-PCR analysis of *ATOH1, BARHL1, OLIG2, CORL2* mRNA levels relative to *GAPDH*(2^-ΔCT^). Data based on five independent differentiation experiments for each condition using F002.1A.13 iPSC line. Mann-Whitney test; error bars represent SEM.

**Supplementary Figure 4.**
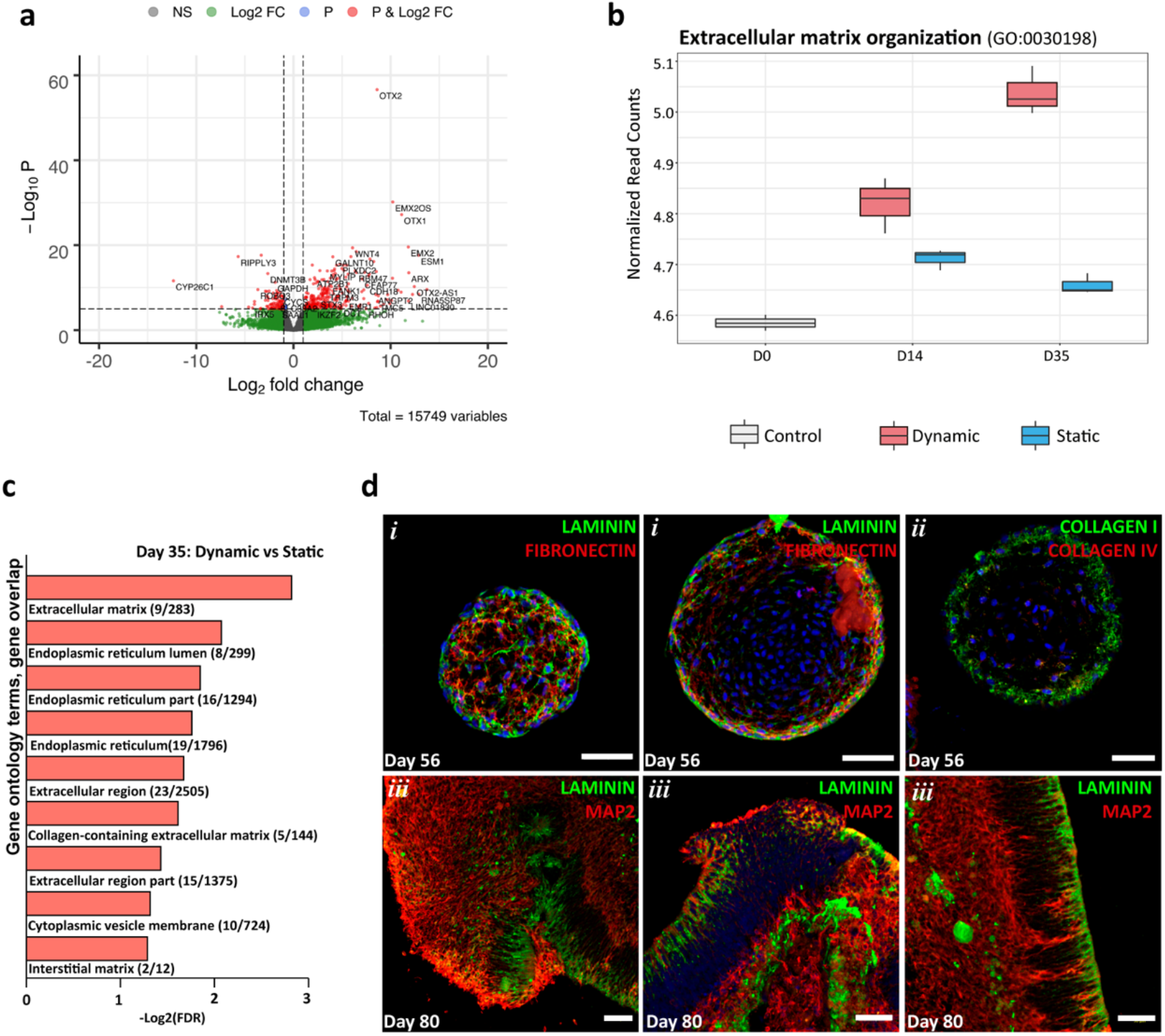
Comparison of effects of dynamic culturing conditions in Extracellular matrix (ECM) remodeling. **a,** Volcano plot of genes identified during cerebellar differentiation of human iPSC using dynamic and static conditions. **b,** Box-plot representing RNA-seq normalized read counts of transcripts annotated in the ECM organization for control (day 0), dynamic and static conditions at indicated time-points of differentiation. **c,** Top gene ontology (GO) cellular component terms identified for the differentially up-regulated genes (Log2 FC > 2 and adjusted p-value < 0.05) of dynamic versus static conditions at day 35. **d,** Immunofluorescence staining for different markers of ECM components and MAP2 on days 56 and 80 of differentiation. Scale bars, 50μm.

**Supplementary Figure 5.**
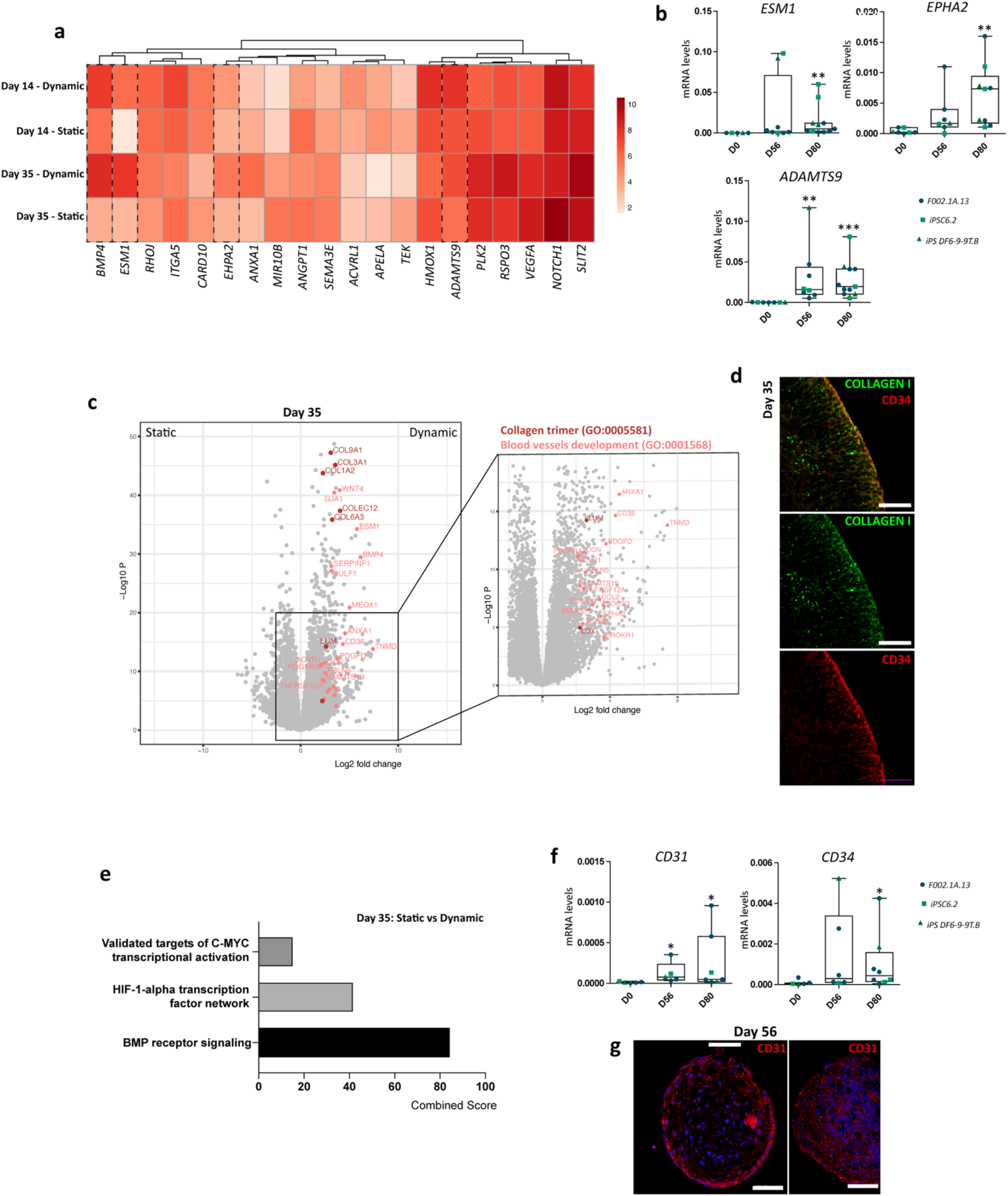
Comparison of effects of dynamic culturing conditions in angiogenesis processes. **a,** Heat map highlighting the top 20 differentially expressed genes related with the sprouting angiogenesis process (GO:0002040) for dynamic and static conditions at days 14 and 35 of differentiation. **b,** qRT-PCR analyses of aggregates derived from F002.1A. 13 iPSC line for genes involved in the sprouting angiogenesis process. Diagrams depict mRNA expression levels (2^-ΔCT^) relative to GAPDH based on three independent differentiation experiments for each condition. One-way ANOVA (Kruskal-Wallis test), *p<0.05, **p<0.01, ***p<0.001; error bars represent SEM. **c,** Volcano Plot of differentially expressed genes between dynamic and static conditions showing significantly up-regulated transcripts in dynamic conditions (Log2 FC > 2 and adjusted p-value < 0.05) related with collagen trimer (GO:0005581) and blood vessels development (GO:0001568) processes. **d,** Immunostaining analyses for COLLAGEN I and CD34 at day 35 in dynamic conditions. Scale bars, 50μm. **e,** NCI-Nature pathways identified for the top 100 genes differentially down-regulated genes (Log2 FC > 2 and adjusted p-value < 0.05) of dynamic versus static condition at day 35. **f,** qRT-PCR analyses of aggregates derived from F002.1A.13 iPSC line for *CD31* and *CD34*. Diagrams depict mRNA expression levels (2^-ΔCT^) relative to GAPDH based on three independent differentiation experiments for each condition. One-way ANOVA (Kruskal-Wallis test), *p<0.05; error bars represent SEM. **f,** Immunostaining analyses for CD31 at day 56 in dynamic conditions. Scale bars, 50μm.

